# Selective Liposomal Transport Through Blood Brain Barrier Disruption in Ischaemic Stroke Reveals Two Distinct Therapeutic Opportunities

**DOI:** 10.1101/570234

**Authors:** Zahraa S. Al-Ahmady, Dhifaf Jasim, Sabahuddin Syed Ahmad, Raymond Wong, Michael Haley, Graham Coutts, Ingo Schiessl, Stuart M. Allan, Kostas Kostarelos

**Affiliations:** Pharmacology Department, School of Science and Technology, Nottingham Trent University, Nottingham, NG11 8NS; Nanomedicine Lab, Faculty of Biology, Medicine and Health, AV Hill Building, Manchester, M13 9PT, United Kingdom.; Division of Neuroscience & Experimental Psychology, Faculty of Biology, Medicine and Heath, University of Manchester, United Kingdom.

## Abstract

The development of new therapies for stroke continues to face repeated translational failures. Brain endothelial cells form paracellular and transcellular barriers to many blood-borne therapies and the development of efficient delivery strategies is highly warranted. Here, in a mouse model of stroke, we show selective recruitment of clinically used liposomes into the ischaemic brain that correlates with biphasic blood brain barrier (BBB) breakdown. Intravenous administration of liposomes into mice exposed to transient middle cerebral artery occlusion took place at early (0.5h and 4h) and delayed (24h and 48h) timepoints, covering different phases of BBB disruption after stroke. Using a combination of *in vivo* real-time imaging and histological analysis we show that selective liposomal brain accumulation coincides with biphasic enhancement in transcellular transport followed by a delayed impairment to the paracellular barrier. This process precedes neurological damage in the acute phase and maintains long-term liposomal co-localisation within the neurovascular unit, which could have great potential for neuroprotection. Levels of liposomal uptake by glial cells are similarly selectively enhanced in the ischaemic region late after experimental stroke (2-3 days), highlighting their potential for blocking delayed inflammatory responses or shifting the polarization of microglia/macrophages towards brain repair.

These findings demonstrate the capability of liposomes to maximise selective translocation into the brain after stroke and identify for the first time two windows for therapeutic manipulation. This emphasizes the benefits of selective drug delivery for efficient tailoring of new stroke treatments.

## Introduction

Stroke is a devastating neurological condition and a leading cause of death and disability worldwide, yet treatment options are extremely limited and thus represent an area of unmet clinical need [1, 2]. At present, restoration of blood flow with thrombolysis and/or thrombectomy are the only licensed treatments for ischaemic stroke, however these options can only be administered up to 4.5h post-stroke, benefitting only a minority of patients.[3] While reperfusion strategies are effective in opening up occluded cerebral vessels in some patients, there are currently no approved treatments for the myriad of damaging pathological processes that persist in the brain long after the acute stage such as oxidative stress and inflammation.[4] Therefore, targeting these downstream pathophysiological processes could hold great therapeutic potential. However considerable research effort has been invested over the last 30 years into the development of novel neuroprotective treatments, with a lack of success. [5]. There are many possible explanations for this translational failure, but insufficient concentrations of drug that reach the intended target area is likely a major factor [6]. Another important problem is the time window between the onset of stroke and treatment initiation which has been frequently wider in clinical trials compared to successful experimental stroke studies [4]. Therefore, developing new technologies that can circumvent inefficient brain delivery and/or unfavourable distribution and safety profiles would lend new prospect to already existing therapeutics.

In normal conditions, brain endothelial cells (BECs), through tightly regulated transcellular transport and tight junction proteins[7], are the primary regulators for the entry of blood-borne molecules into the brain. During stroke there is strong evidence from preclinical[8-10] and clinical[11, 12] studies that blood brain barrier (BBB) integrity is compromised. Ischaemic conditions affecting the brain tissue alter the rate and the extent of BEC transcellular transport and change the expression levels and localisation of tight junction proteins.[9] Moreover, degradation of extracellular matrix by proteolytic enzymes (such as matrix metalloproteinases) [13, 14], release of inflammatory mediators and infiltration of peripheral blood leukocytes [15] have all been proposed to contribute to BBB hyperpermeability after ischaemic stroke. As a result, a biphasic increase in BBB hyperpermeability and uncontrolled entry of molecules to the brain occurs. The exact contribution of transcellular and paracellular pathways to BBB hyperpermeability after ischaemic stroke is a matter of controversy[16]. However, the most accepted model of hyperpermeability is characterised by; a) an early phase (occurring a few hours post stroke) of enhanced transcellular transport mediated by increases in endothelial vesicles termed caveolae, followed by b) a delayed phase of hyperpermeability (~ 2d post-stroke) in which both enhanced transcellular transport and tight junction proteins disassembly contribute to the loss of BBB integrity[9]. This two-phase model is supported by recent evidence in which endothelial tight junctions were labelled with eGFP, allowing the dynamics of tight junction integrity after stroke in mice to be monitored *in vivo* in real time[9]. Similarly, in a rat model of stroke it was shown that BBB opening to macromolecules (transcellular route) precedes permeability to small ions (paracellular route)[17]. Moreover, recent findings in a comorbid rodent model of ischaemic stroke, reported a significant exacerbation to the BBB disruption when combined with other diseases without necessarily altering the underlying sequence of events behind BBB disruption[10]. Consequently, BECs that survive the ischaemic damage, but do not maintain BBB integrity, can in fact worsen the damage to the brain parenchyma and thus accelerate disease progression [18]. On the other hand the disrupted BBB could act as a gate for therapeutic access.[7] This highlights the need for more effective delivery approaches that can selectively and efficiently penetrate areas of BBB hyperpermeability compared to other brain regions where BBB permeability is unaffected.

In the past few decades, nanotechnology-based drug delivery approaches such as liposomes have demonstrated a great potential to improve the pharmacokinetics and biodistribution profile of many small drug molecules. Selective accumulation of liposomes into the disease site, such as a tumour, is mediated by hyperpermeability of endothelial cells and impaired lymphatic drainage compared to other healthy organs. This phenomenon is collectively known as the enhanced permeability and retention (EPR) effect[19], which is behind the clinical use of many liposomal-based medications[20]. Previous studies have shown that endothelial cell hyperpermeability is mediated by one or more of the following pathways; a) fenestrations in the basement membrane[21], b) paracellular transport through gaps between endothelial cells[21], and c) transcellular route mediated by vascular transport (caveolae)[22]. These pathways are in great analogy to BECs structural adaptations after ischaemic stroke and, therefore, highlight the possibility that liposomes might be equally effective in the treatment of stroke by providing selective and enhanced drug delivery to the ischaemic brain. In this respect there are a few promising examples of liposomal treatment of ischaemic stroke reported recently[23–28]. However, the focus of those studies is restricted to the acute phase (<1-3h) after ischaemic reperfusion and they did not fully address longer-term pathological consequences and the practicality of clinical translation. Given that the main aim of neuroprotective treatment of stroke is to expand the therapeutic window compared to thrombolytic therapies, an in depth understanding of the window of BBB opening after stroke is required to fully achieve the potential of liposomes as a drug delivery approach.

The aim of this study was to fully interrogate the validity of utilizing liposomes to maximise drug delivery in stroke. Multiple quantitative and qualitative techniques were employed to establish the time window and mechanism of liposomal accumulation into the brain after experimental stroke in mice. Early and delayed accumulation of liposomes into the brain were evaluated and time points that gave rise to key brain accumulation are highlighted for future therapeutic evaluation.

## Results

In order to establish the validity of liposomes for selective drug delivery into the ischaemic brain, we have applied *in vivo* and *ex-vivo* imaging techniques to study their recruitment into the brain after stroke. A liposomal formulation based on the composition of clinically used liposome (Doxil^®^, HSPC:CHOL:DSPE-PEG_2000_) was selected, as this has proved to have excellent blood circulation time and very good drug retention capability once inside the body[29] (For characterisation data please see Figure S1). Selective recruitment of liposomes into the brain was studied in a preclinical model of middle cerebral artery occlusion (MCAo) followed by reperfusion.

### Selective liposomal accumulation into the ischaemic brain correlated with biphasic BBB breakdown induced by stroke

Liposomal accumulation into the ischaemic brain was first studied shortly after intravenous (I.V) administration of fluorescently-labelled liposomes (DiI-Lp). Each group of mice received a single I.V injection of DiI-Lp. The time points of injections were carefully selected to cover the two-phases of BBB damage after ischaemic stroke as previously reported [9]. IVIS Lumina II imaging (Figure 1A) confirmed the selective accumulation of liposomes into the ischaemic left side of the brain as early as 2h after DiI-Lp I.V administration compared to minimum detection from the contralateral side of the brain (right) and healthy mice (naïve mice injected with DiI-Lp). Quantification of the total fluorescent signal of liposomes in the brain indicated that the accumulation of liposomes significantly increased when injected at 0.5h or 48h following MCAo and reperfusion compared to healthy mice injected with DiI-Lp (Figure 1B). This suggested that liposomal brain accumulation induced by stroke is correlated with the biphasic BBB breakdown in agreement with previous reports.[9, 14, 30, 31] Histological analysis of the brain tissue confirmed that the selective liposomal accumulation into the ischaemic area has the same distribution to BBB disruption (Figure 1D), as seen by infiltration of endogenous immunoglobulin (IgG).

**Figure 1:**
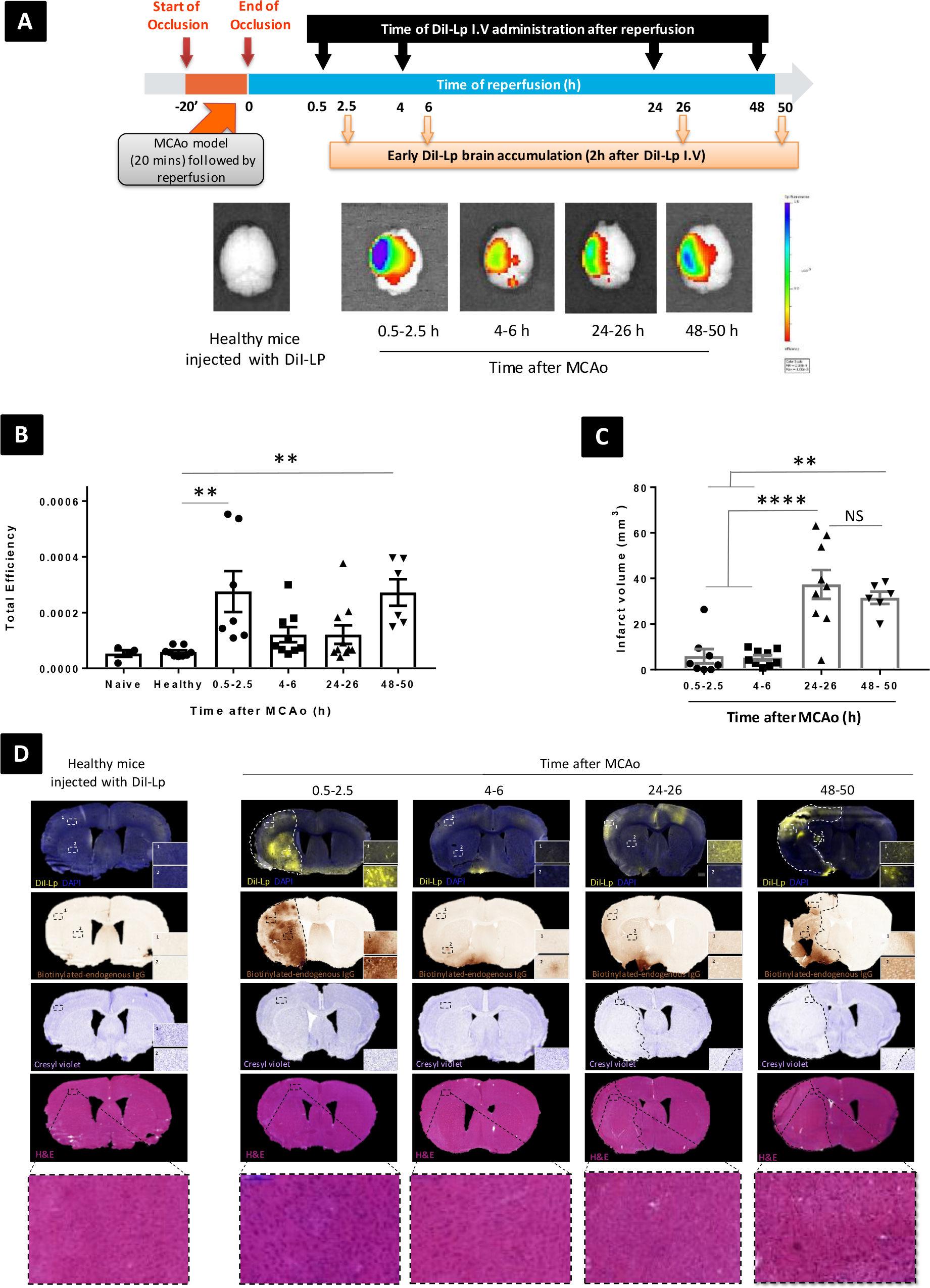
Selective liposomal accumulation into the ischaemic brain shortly after Lp-DiI I.V administration. Selective recruitments of DiI-Lp into the ischaemic region of the brain in MCAo model of stroke were studies by (A) IVIS Lumina II imaging system showing the selective accumulation of liposomes into the ischaemic left side of brain as early as 2h after DiI-Lp I.V administration. Minimum DiI-Lp signal was detected from the contralateral side of the brain (right) and healthy mice (naive mice injected with liposomes). (B) Quantification of the fluorescent single of DiI-Lp in the brain by IVIS Lumina imaging software was performed by drawing a region of interest (ROI) that covers the whole brain and expressed as total efficiency. Colour scale of epi-fluorescent signal range from min= 2.00^−4^ (red) to max= 1.00^−3^ (blue). A bi-phasic recruitment of DiI-Lp into the ischaemic area was observed showing significant increase when injected 0.5h or 48h following MCAo and reperfusion. (C) Quantification of the volume of ischaemic damage was performed on representative sections taken at eight defined coronal levels. (D) Brain sections from healthy mice and mice after MCAo showing; maximum selective recruitment of DiI-Lp into ischaemic left side of the brain (marked by dashed white lines at 0.5-2.5h and 48-50h time point), compared to less liposomal accumulation at 4-6h and 24-26h groups and no liposomes accumulation into the healthy brain. Accumulation of liposomes into the ischaemic brain have a similar distribution to the endogenous IgG leakage into the brain (outline by dashed black lines) that is used as indication of BBB disruption. Cresyl violet staining of brain sections represents the extent of brain damage induced at different time points following 20min MCAo. In most cases moderate to extensive damage of both cortical and subcortical regions was observed at 1-2d post MCAo. Cerebral accumulation of DiI-Lp given at the early phase after reperfusion (0.5h & 4h) were observed before clear histological evidence of neuronal damage was observed as indicated from the cresyl violet stain. H&E images confirmed that liposomal accumulation was associated with rare instances of vessel lumen collapse. Inset images represent 40 x magnifications for DiI-Lp and 20x magnification for IgG, cresyl violet and H&E. n=5-7 in each group.

### Liposomal brain accumulation in the acute phase post ischaemic stroke preceded neurological damage

Cerebral accumulation of DiI-Lp injected at the early phase after MCAo and reperfusion (0.5-2.5h & 4-6h) were observed as early as 2h after I.V administration, before any histological evidence of neuronal cell death (Figure 1C&D). This early accumulation was associated with rare instances of vessel lumen collapse as seen by H&E staining (Figure 1D), but in the absence of any active bleeding process.

### Liposomes maintained selective accumulation in the ischaemic area of the brain 24h after I.V administration

Cerebral accumulation of DiI-liposomes given 0.5h and 48h after reperfusion was still observed in the ischaemic region 24h after their I.V injection and maintained similar distribution to the areas of BBB damage and infarct as seen with IgG infiltration and Cresyl violet stain respectively (Figure 2). Liposomal accumulation at the time windows of maximum selective recruitment into the brain (0.5h & 48h) were also studied in real time by SPECT-CT imaging (Figure 3A) using ^111^In-DTPA-liposomes (^111^In-Lp). Characterisation data of ^111^In-Lp and radiolabelling efficiency are explained in Figure S2. During real time SPECT/CT imaging the selective recruitment of liposomes in the brain became more apparent 24h after injection since immediately after injection the high level of liposomes in the blood gave rise to equal signal from both sides of the brain (Figure 3B). Quantification of ^111^In-Lp accumulation 24h after I.V administration confirmed significant increase in ipsilateral accumulation of ^111^In-Lp when injected at 0.5h or 48h after MCAo (Figure 3E). On the contrary no significant difference in ^111^In-Lp levels was detected in the contralateral region (Figure 3F). Measurements of ^111^In-Lp in the CSF (Figure 3G) indicated no significant increase in liposomal clearance into the CSF after MCAo, which explained the persistence of liposomes in the ischaemic area even 1d after administration.

**Figure 2:**
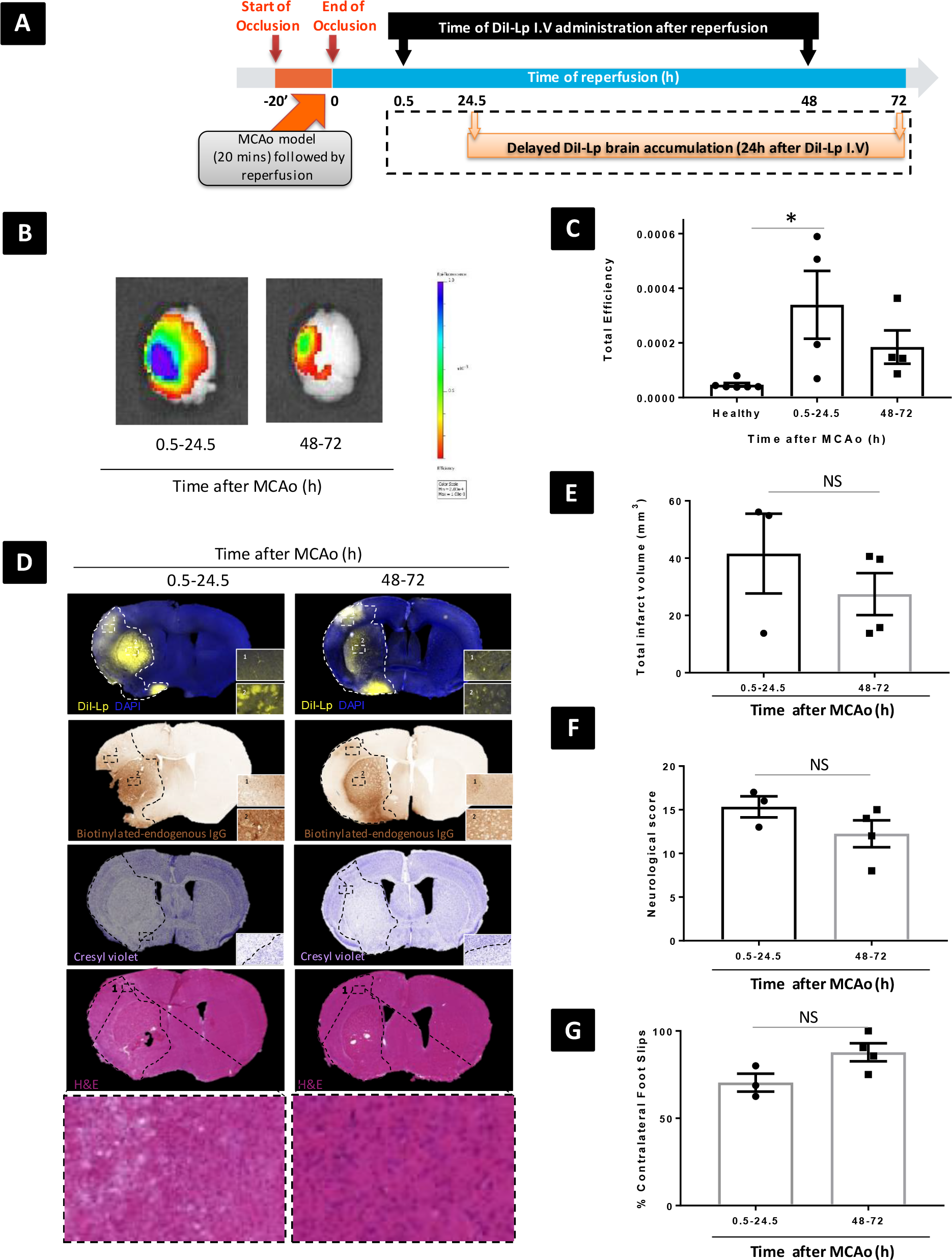
Selective liposomal accumulation into ischaemic brain 24h after I.V administration. (A) Schematic presentation of the experimental time points used for DiI-Lp I.V administration following 20min MCAo and reperfusion. The time points of DiL-Lp intravenous injection illustrated the separate groups studied each received single injection. Selective recruitments of DiI-Lp into the ischaemic region of the brain in MCAo model of stroke were studies by (B) IVIS Lumina II imaging system showed that selective accumulation of liposomes into the ischaemic (left side) persists 24h after DiI-Lp I.V administration. (C) Quantification of the total fluorescent single of DiI-Lp in the brain 24h after injection indicated that there was no change in liposomal accumulation when injected 0.5h after MCAo, whereas reduced total liposomal level was observed when injected 48h after MCAo. Colour scale of epi-fluorescent signal range from min= 2.00^−4^ (red) to max= 1.00^−3^ (blue). A bi-phasic recruitment of DiI-Lp into the ischaemic area was observed showing significant increase when injected 0.5h or 48h following MCAo and reperfusion. (D) Cerebral accumulations of DiI-liposomes administered 0.5h & 48h after reperfusion were still observed in the ischaemic region 24h afterwards and observed co-localised with BBB damage (endogenous IgG) and infarct area (Cresyl violet) with no clear evidence of vessel lumen collapse (H&E). Inset images represent 40 x magnifications for DiI-Lp and 20x magnification for IgG, cresyl violet and H&E. Primary brain injury after MCAo was confirmed by (E) Quantification of volume of ischaemic damage on representative sections taken at eight defined coronal levels of brain tissues stained with cresyl violet, (F) measuring focal deficit scoring (0-28) and (G) foot-fault test.

**Figure 3:**
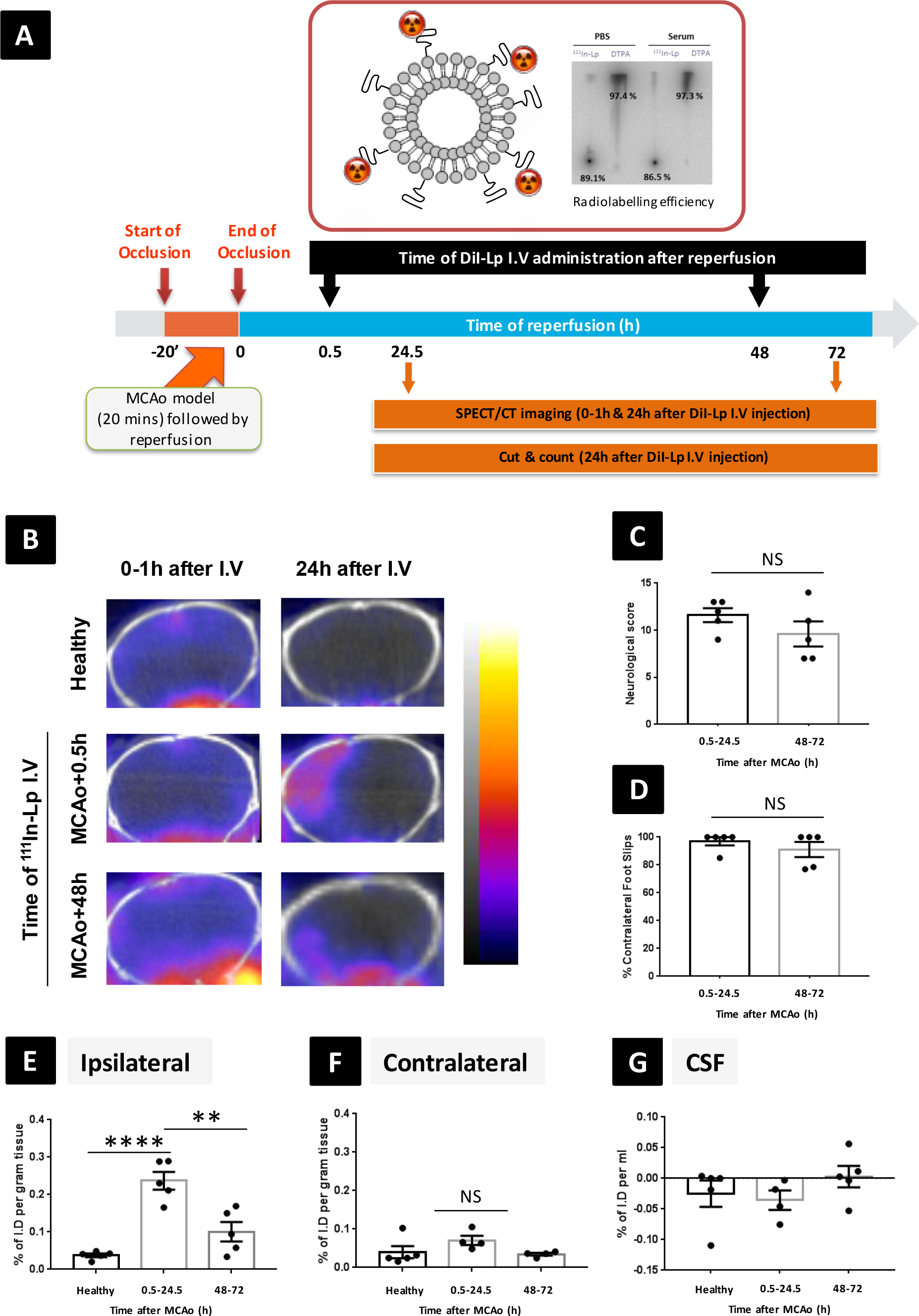
Quantification of liposomes accumulation into the brain after MCAo. (A) Schematic presentation of experimental design and ^111^In-Lp radiolabelling efficiency. The time points represent separate groups each received single injection of ^111^In-Lp intravenously. TLC Data indicated > 85% radiolabelling efficiency with excellent serum stability. (B) Representative SPECT/CT imaging confirmed the selective accumulation of the liposomes into the ipsilateral side of the brain (left) compared to contralateral side. In the absence of ischaemic stroke, no accumulation of ^111^In-Lp was detected. Assessment of (C) neurological focal deficit scoring (0-28) and (D) foot-fault test showed no significant differences between 0.5h (MCAo +1d) and 48h groups (MCAo +3d). Quantification of ^111^In-Lp accumulation into (E) ipsilateral brain side and (F) contralateral brain side 24h after I.V administration. Values are expressed as % of I.D ± SEM per gram brain tissue. Data confirmed significant increase in selective brain liposomal level when injected at 0.5h after MCAo and to a less extent when given 48h after MCAo. (G) Detection of ^111^In-Lp in the CSF after MCAo indicated no significant differences in CSF liposomal level compared to healthy control group.

### Biphasic increase in liposomal brain accumulation after ischaemic stroke connected with enhanced transcellular transport

To gain an in depth understanding of the mechanisms of selective liposomal recruitment into the brain after stroke, we investigated the molecular pathways (transcellular vs paracellular) that could be involved. Experimental MCAo model leads to a biphasic increase in transcellular transport across the BBB in the early and delayed phases after ischaemia (Figure 4A). However, it is unclear if enhanced transcellular transport (endothelial caveolae) is connected to the biphasic increase in liposomal brain accumulation after ischaemic stroke. Caveolae are invaginations of plasma membrane that play an important role in the transcellular transport of small molecules such as cholesterol and albumin [32]. Among the caveolins family, caveolin-1 (Cav-1) is known as an important regulator of caveolin mediated transport. The exact contribution of Cav-1 to BBB hyperpermeability is not clear, though recent findings indicate enhanced Cav-1 expression by BECs and increased numbers of transcellular vesicles shortly after ischaemic insult and before tight junction (TJ) disassembly [9, 33, 34]. Moreover, Knowland et al confirmed that this effect is in fact biphasic and contributes to both early and late BBB disruption [9]. Consistent with previous reports [35, 36], TEM images of brain tissues in the acute (MCAo+2.5h) and late (MCAo+50h) phases after stroke (Figure 4B) confirmed ultrastructural changes in endothelial cells compared to healthy brains. Increases in caveolae numbers were clearly evident in the cytoplasm of BECs at both time points. Enlargement of the vesicles and ultrastructural changes to endothelial cell TJs (protrusions and change in morphology) were also observed at the late but not the early time point after MCAo. These observations were further confirmed by immunofluorescence labelling of endothelial CD31 and Cav-1 markers. Analysis of the brain tissues 2h after intravenous administration of DiI-Lp revealed a biphasic increase in Cav-1 expression from 0.5h (MCAo+2.5h) and 48h groups (MCAo+50h) that co-localised with the areas of DiI-Lp leakage into the ischaemic brain (Figure 4C&D). These observations were consistent from both ipsilateral striatum (Figure 4C) and cortex (Figure S3). On the contrary, Cav-1 expression and accumulation of DiI-Lp in the brain were minimum from MCAo groups injected at 4h & 24h post stroke, healthy group and contralateral sides (Figure S5 & Figure S6). Evaluation of DiI-Lp localisation 24h following I.V at 0.5h and 48h to MCAo mice indicated that substantial numbers of liposomes maintain their co-localisation in the neurovascular unit (NVU) in areas of enhanced Cav-1 expression (Figure 4E & Figure S4).

**Figure 4:**
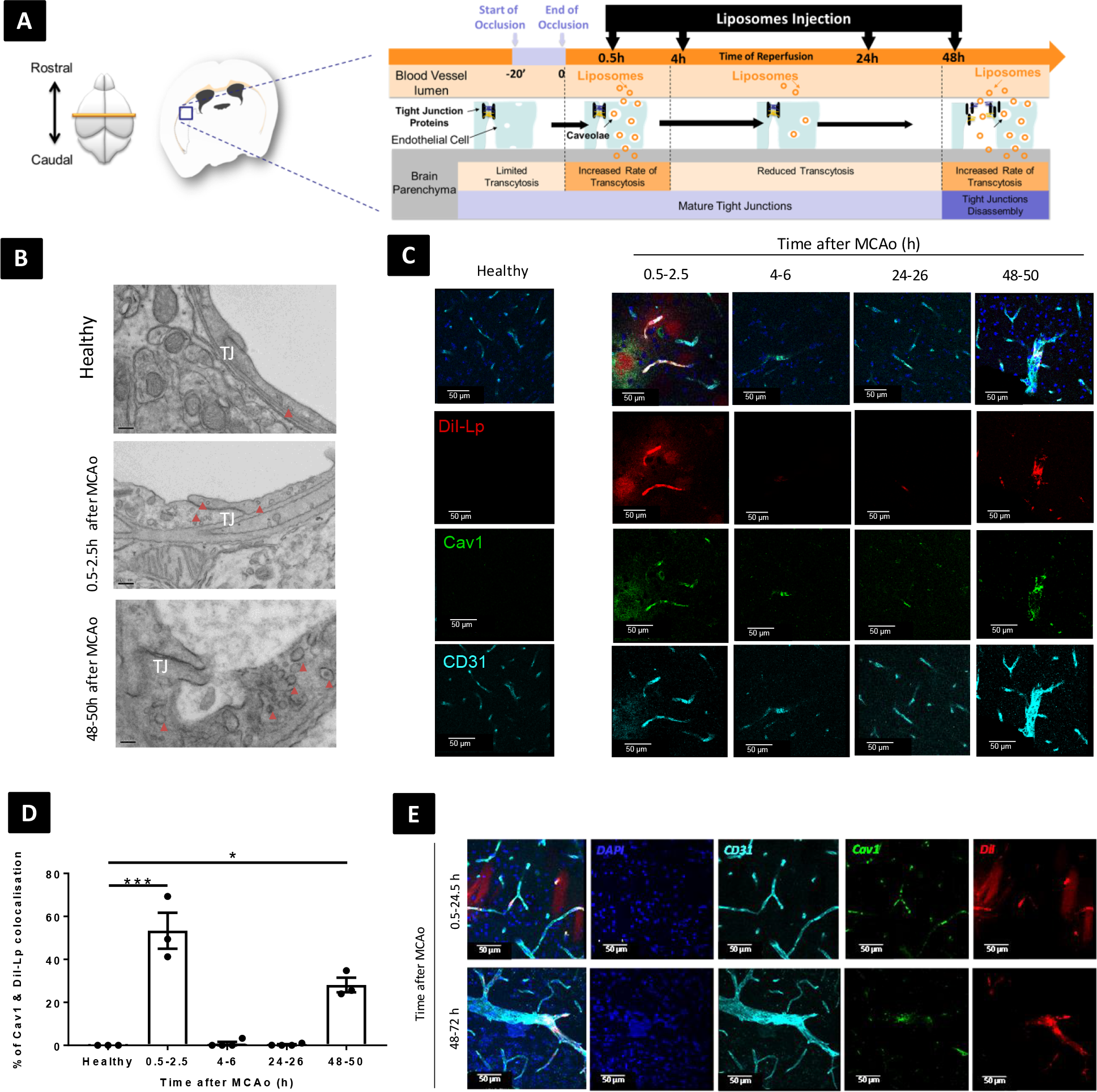
Mechanism of selective liposomal accumulation into ischaemic brain. (A) Schematic presentation of the experimental plan and the time frame for DiI-Lp intravenous administration after 20min of MCAo model in mice. The time points represent separate groups each received single injection of DiI-Lp intravenously. The two possible mechanisms of selective liposome recruitment into the ischaemic brain are elucidated. (B) Representative TEM images confirmed ultrastructural changes in endothelial cells after MCAo compared to healthy mice. Amplification of transcytotic vesicles number was clearly evident at 0.5-2.5h and 48-50h after MCAo. Enlargement of the vesicles and ultrastructural changes to endothelial cells TJs (protrusions and change in morphology) were observed at late but not early time points after MCAo. (C) Representative images demonstrating immunofluorescence labelling of endothelial CD31 (cyan) and Cav-1 (green) markers in the striatum at −0.58mm from the bregma. Analysis was performed 2h after I.V injection of DiI-Lp for each group. A biphasic increase in Cav-1 immunofluorescence was observed for 0.5-2.5h and 48-50h groups that co-localises with the areas of DiI-Lp leakage into the ischaemic brain. On the contrary, Cav-1 expression and accumulation of DiI-Lp in the brain were minimum for 4-6h and 24-26h groups. (D) Quantitative analysis of Cav1 co-localisation with DiI-Lp 2h after I.V injection confirmed the biphasic pattern observed from confocal images. (E) Evaluation of DiI-Lp leakage into the ischaemic brain 24h following I.V into MCAo mice at 0.5h and 48h following reperfusion. Co-localisation of DiI-Lp with areas of enhanced Cav-1 expression were detected.

### Delayed liposomal brain accumulation after ischaemic stroke co-localised with regions of impaired paracellular barrier

In addition to enhanced transcellular transport, compromised paracellular barrier by virtue of TJ proteins disassembly leads to BBB disruption after ischaemic stroke. Up to 30% of tight junction strands were reported to be open 48–58h after MCAo and the opening of these gaps increased progressively after stroke, reaching gaps of 0.2-1.2 µm.[9] Since the size of TJ opening is far bigger than the hydrodynamic diameter of liposomes tested in this study (120-130nm) it is feasible that the delayed phase of liposomal accumulation is facilitated by the paracellular route. To test this, mice were implanted with a cranial window underwent MCAo surgery and received DiI-Lp intravenously at 0.5h or 48h after MCAo. Together with DiI-Lp injections, mice also received I.V injections of fluorescent tracers for paracellular pathway (dextran 3k) and transcellular pathway (albumin-Alexa488). To examine the dynamic accumulation of liposomes into the ischaemic brain and correlate that with structural abnormalities of BECs, we subjected MCAo mice to *in vivo* multiphoton imaging recording at 30 min and 2 h after DiI-Lp and tracers injections. These time points were selected to match our previous data sets and offers the advantage of having both time points in the same animal.

Dextran is normally excluded from the brain parenchyma by an intact TJ, however, its extravasation outside the blood vessels increased in the late phase after ischaemic stroke progression (Figure 5). This extravasation became apparent at 48-50h after MCAo and reperfusion, correlating with the TJ disassembly. In contrast, albumin uptake by BECs, which was reported to be through caveolae mediated endocytosis[37], increased both in the acute (0.5-2.5h) and late (48-50h) phases after MCAo, corroborating our previous observations from TEM and IHC analysis. These data were validated *ex vivo* and on brain sections, using IVIS (Figure 5D&E).

**Figure 5:**
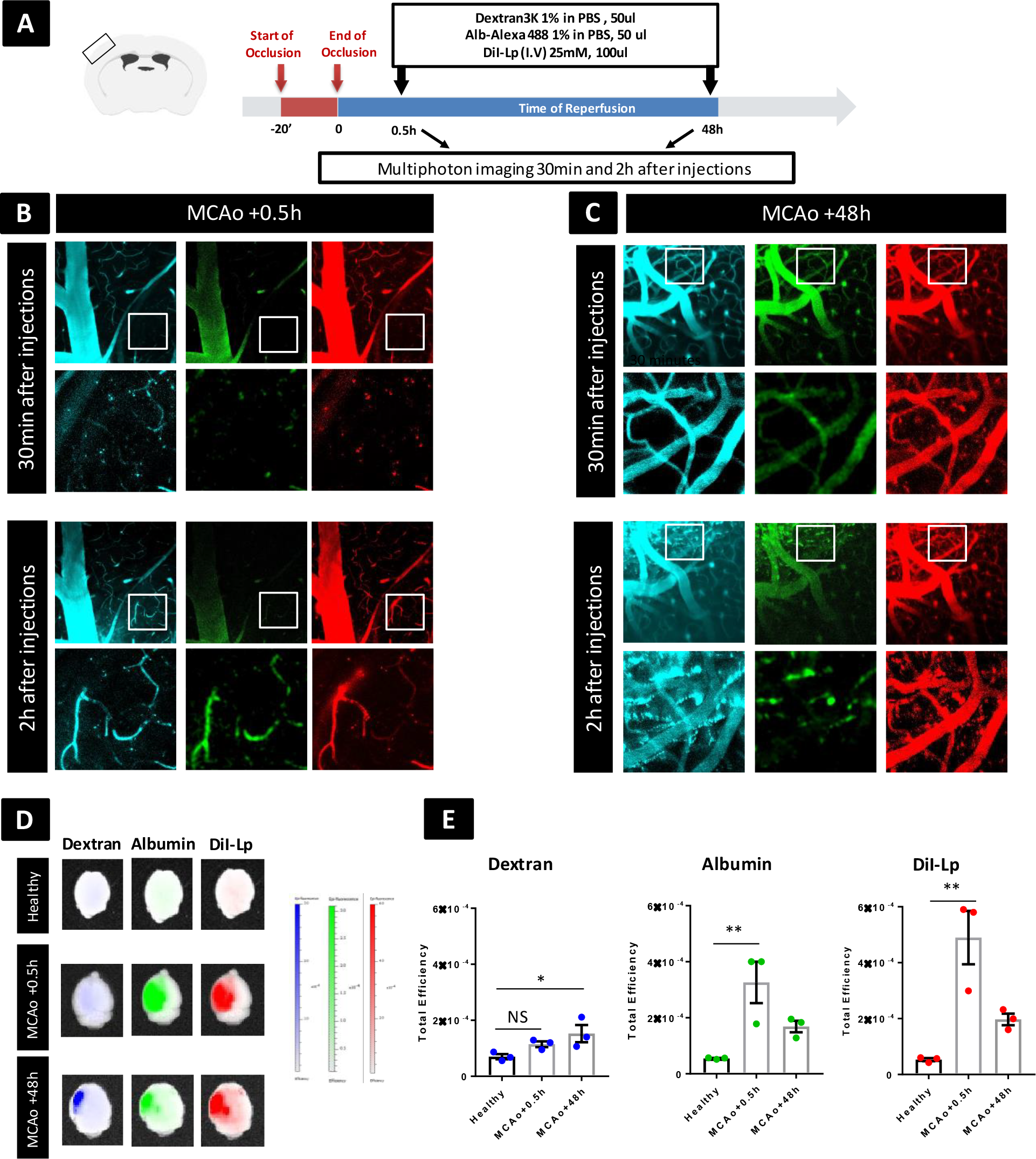
Co-localisation of liposomal accumulation into ischemic brain with transcellular and paracellular markers. (A) schematic presentation of study design for multiphoton imaging. In this study each mouse received I.V injections of DiI Lp (25mM, 100µL), cascade blue dextran 3K (50µL) and albumin-Alexa488 (50µL) either in the early phase (MCAo+0.5h) or delayed phase (MCAo+48h) after stroke. (B&C) Mice imaged with SP8 upright multiphoton microscope using 25x lense with water immersion. Imaging performed 30 min and 2h after injections represented the distribution of dextran (cyan), albumin (green) and DiI-Lp (red) into the ipsilateral brain. The wide view images in (B) and (C) are 442 µm wide and the 130 µm wide ROI outlined by the white square is shown below each image. (D) At the end of the experiment mice were perfused with saline and fixed with PFA and the brains were collected and imaged with IVIS Lumina II imaging system. (E) Quantification of the fluorescent single of dextran, albumin and DiI-Lp in the brain by IVIS Lumina imaging software was performed by drawing ROI that covers the whole brain and the data expressed as total efficiency. Colour scale of epi-fluorescent signal range from min= 1.00^−5^ to max=3.00^−4^ for dextran and albumin and from min= 1.00^−5^ to max=6.00^−4^ for DiI-Lp.

Overall, our data suggest that biphasic upregulation of transcellular pathway followed by delayed TJ disassembly are the drive for the selective liposomal accumulation into the ischaemic side of the brain. Although Cav-1 is known to be essential for transcellular transport of albumin through BECs, recent observations showed that albumin uptake could still be enhanced in Cav-1 deficient mice, though to a lesser extent, which imply the contribution of other Cav1-independent pathways[9].

### Selective targeting of ischaemic brain with liposomes offers time-dependent tailoring of stroke treatment

Immunobiological analysis described above indicated that liposomes administered in the acute (0.5h) and late (48h) phases after stroke co-localised with the NVU. Since the NVU plays a critical role in the progression of ischaemic damage after stroke, selective targeting of the NVU with liposomes, particularly in the acute phase, could hold great potential as a therapeutic strategy by limiting the deleterious effects of BBB disruption after stroke.

In addition to the NVU, co-localization of the liposomes with other cellular compartments of the brain parenchyma was tested. Immunostaining for ionised calcium binding adaptor molecule 1 (Iba-1), which is exclusively expressed by macrophages and microglia [38] and glial fibrillary acifdic protein (GFAP), expressed by astrocytes, was performed (Figure 6). The percentage of DiI-Lp uptake by Iba1+ cells and GFAP+ cells was quantified 24h after I.V DiI-Lp. We found that a significant fraction (40-50%) of DiI-Lp co-localised with Iba1+ cells when administered 2 days after MCAo compared to <3-4% when injected in the acute phase, at 0.5h after MCAo (Figure 6F). Resting microglia dynamically monitor the brain microenvironment [39], and become activated when sensing damaging signals [40], changing morphology from long, ramified protrusions to shorter, hyper-ramified branches.[40] Our results demonstrate DiI-Lp uptake by both resting and activated microglia, though it is enhanced in the latter (Figure 6G). Minimum uptake of DiI-Lp by astrocytes was observed at both times tested (Figure 6H). No specific uptake of DiI-Lp by neurons was detected at the time points tested, while occasional co-localisation with MBP+ cells (oligodendrocytes) was observed (Figure S7 & Figure S8).

**Figure 6:**
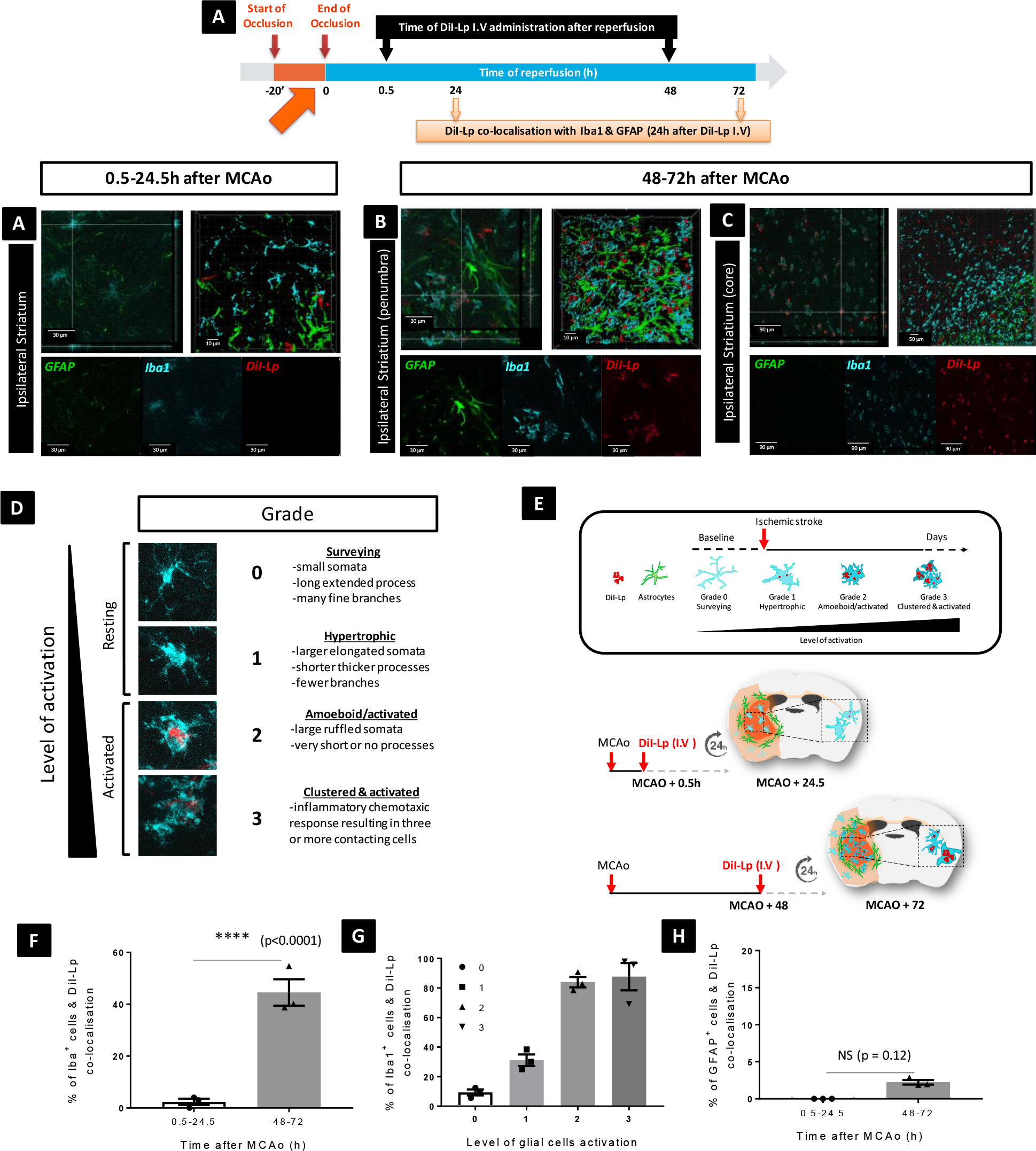
Evaluation of DiI-Lp co-localisation with microglia and astrocytes. Representative confocal images showing co-localisation of DiI-Lp with of Iba1 (activated microglial marker) and GFAP (astrocytes marker). Brain sections were analysed 24h following the injection of DiI-Lp intravenously into MCAo mice in the; (A) early phase (MCAo+0.5h) and (B&C) delayed phase (MCAo+48h) after reperfusion. (D) Scoring system of microglial activation showing representative images of the different stages of microglial cells activation on a scale from 0-3. (E) Schematic summary of liposomal co-localisation with microglia and astrocytes. (F) Quantification of DiI-Lp co-localisations indicated significant uptake of DiI-Lp by microglia when injected 48h after MCAo. (G) The correlations of DiI-Lp uptake by microglia with the different stages of microglial activation after MCAo. Significant uptake of liposomes by activated microglial was observed compared to minimum uptake by resting microglia (H) No significant uptake of DiI-Lp into astrocytes was detected.

## Discussion

Stroke-induced BBB disruption results in a biphasic enhanced entry of blood borne molecules to the brain. Despite the obvious benefits this can offer to enhance drug delivery to the ischaemic brain, this area has been largely overlooked. Understanding the link between the response of BECs *in vivo* to ischaemic injury and selective drug delivery to the brain is crucial to the development of timely therapies that may halt disease progression.

Recent studies using nanoparticles-based delivery systems demonstrated the possibility to selectively target the lesion area after experimental stroke. This effect was only demonstrated when those nanoparticles injected I.V just before reperfusion or 1-3 h afterward [41, 42]. Although these studies suggested that the increase in the permeability of the BECs is the drive for this selective accumulation, no direct correlation to BBB damage was reported. Moreover, the exact mechanisms by which the selective brain localization happens are largely unknown and require in depth studies. Therefore, in this study we systematically interrogated the benefit of BBB damage to enhance drug delivery by analysing the brain accumulation and localisation of I.V liposomes injected at different time points after experimental stroke to cover the biphasic increase in BBB permeability. The most acceptable model of BBB response to ischemia involves a stepwise activation of two distinctive pathways, starting with enhanced BECs transcellular transport early after reperfusion followed by a second delayed phase (~48h post stroke) of enhanced transcellular transport and major disassembly to TJ protein complexes [9, 43].Here we demonstrate for the first time that biphasic BBB hyperpermeability after stroke correlates with selective enhancement of liposomes accumulation into the ischaemic brain. More specifically we identified two distinct windows for maximum localisation of liposomes into the brain; an early phase (0.5h after stroke) that agrees with previous studies, and a delayed phase (48h after stroke) which has not been reported before. These observations were based on *in vivo* (SPECT/CT and multiphoton imaging) and ex-vivo (IVIS) imaging techniques and were further confirmed by histological analysis. We have demonstrated that enhanced transcellular transport is the main mediator for increase liposomal brain accumulation in the early phase (0.5h). Whereas the reduction in the level of these vesicles in the time between 4h and 24h after stroke is behind the minimal accumulation of liposomes observed at those time points. This agrees with previous studies that reported limited liposomal translocation or therapeutic effect from delivery systems administered around those times [41, 42]. Our data also reveals a second window for liposomal entry into the brain (~48h after stroke) in which both transcellular and paracellular pathways contribute to the selective localisation of liposomes.

The reasons behind the initial increase in transcellular transport through BECs after ischemia are not well defined yet. One possible explanation, correlates with the early migration of pericytes away from the BECs that is triggered after basement membrane dissolution [44] [45]. Pericytes are known to secrete inhibitory signals that reduce the rate of transcellular transport through BECs. Thus, the loss of those inhibitory signals can lead to the initial rise in transcellular transport [46] [47]. Caveolae are the intermediaries of this transcellular pathway and contain receptors to molecules that must cross the BBB [48], However, it is unclear if this increase has any benefit the to NVU at this stage after stroke.[9] The late increase in BBB hyperpermeability is more linked to TJ disassembly triggered by matrix metalloproteinases (MMPs), secreted by inflammatory cells, and angiogenic growth factors such as vascular endothelial growth factor and & nitric oxide synthase [9, 15]. This comes in parallel with a second phase of increase in BECs transcytosis.[9] Although previous studies reported reduced level of TJ proteins (Claudin5, Occludin, and Zona Occludens-1) in the first few hours after stroke [31], ultrastructural analysis of TJ during the evolution of ischaemic stroke are not persuasive that this reduction is responsible for the early BBB breakdown [34, 49]. Based on the studies mentioned above, it is suggested that active disassembly of TJ proteins occur in the late phase of reperfusion, when angiogenesis of CNS vessels begins [50]. This is also in agreement with our data that demonstrated TJ modification only 48h after MCAo. This was confirmed with TEM imaging of brain sections and multiphoton and IVIS imaging of dextran 3k extravasation into the brain.

Identification of the exact therapeutic benefit of this selective brain delivery approach needs substantially more experimental work. Here we provide a clear evidence that selective accumulation of liposomes into the ischemic hemisphere precedes neuronal damage which make them ideal for neuroprotection. Few neuroprotective drugs have been tested in liposomal formulation injected at reperfusion or shortly after such as erythropoietin[26], tacrolimus [41] and CDP-choline [28]. When compared to free drugs, promising therapeutic effects were reported including reduced cellular apoptosis, brain infarct volume and improved motor function.

Beside protecting the brain tissue, NVU itself can be a therapeutic target. It is well known that disruption of the BBB markedly influences the pathogenesis of stroke. Moreover, both systemic and cerebrovascular inflammation contribute to further BBB disruption and can alter stroke prognosis. This effect is both neutrophil and MMPs dependent [51]. Neutrophils are generally considered the first responders to ischaemic stroke as they infiltrate the brain within the first few hours after the ischaemic insult. Neutrophils have a key role in BBB disruption as they are the source of various proteolytic enzymes in particular MMPs. Consistent with that, recent studies indicated that active MMP2/9 observed in the brain after the initial disruption of BBB permeability, approximately 3h after ischemia [52], and progressively increased afterwards [53]. *In vitro* and *in vivo* studies suggested that inhibition of MMP2/9 minimised BECs permeability and transiently reduced the infarct volume. Our immunobiological analysis indicated substantial co-localisation of liposomes with NVU both in the early and delayed phases after stroke which was maintained even 24h after administration. The therapeutic potential of this has not been tested before. However due to the great influence of NVU on the progression of ischaemic damage after stroke, selective targeting of the NVU with liposomes, could be used to influence neutrophil infiltration and/or for MMPs inhibition in order to mitigate the deleterious effects of BBB disruption after stroke.

Cerebral inflammation is also a key factor in neuronal damage after acute ischaemic injury [54]. This is mainly mediated by the secretion of pro-inflammatory mediators by classically activated M1 microglia and macrophages such as interleukin-1 family which are triggered in response to sterile inflammation [55]. However, recent studies highlighted the dual role of microglia/macrophages in brain disorders. In fact, alternatively activated M2 phenotype are the source of protective and neurotrophic factors that reserve brain function and promote brain functional recovery [56]. Therefore, blocking the inflammatory responses after stroke [57–59] or shifting microglia/macrophage polarisation towards brain repair [60] is another area of therapeutic potential. The present study is the first to identify the intriguing selective uptake of liposomes by microglia in the late phase after stroke and its correlation with microglia activation stage. Therefore, it will be essential to investigate the potential of liposomes to tip the central inflammatory responses after stroke towards brain repair. This would allow cell type specific intervention to be tested in timely manner after stroke.

The therapeutic targets described above are driven by the natural tendency of selective liposomal accumulation into the brain post stroke, however, active targeting of specific cell population in the brain is another area worth of investigation. This concept can be highly relevant for delivering neuroprotective drugs to neurons early on after stroke, as liposomal accumulation in the brain was evident prior to neuronal cell loss. A key advantage liposomes offer compared to other drug delivery approaches, is their ability to encapsulate hydrophilic and hydrophobic molecules. This makes them very attractive for stroke therapy, since the choice of encapsulated therapeutic molecules can be easily tailored to match the therapeutic target in a timely manner.

### Limitations of the results

In this study we have established the potential of utilizing liposomes to achieve selective enhanced translation in ischaemic brain after experimental stroke. We have identified two-windows for maximum accumulation into the brain with two distinct therapeutic targets. What we haven’t showed in this study is the translation of this enhanced accumulation into improved therapeutic activity or functional recovery by encapsulating therapeutic molecules inside the liposomes. However, this study lays the basis for critical design of future therapeutic studies to prove the potential liposomes offer to accelerate the clinical translation of stroke treatments.

## Conclusions

We propose liposomal drug delivery to enhance the translocation of therapeutic molecules into the brain by taking advantage of BBB disruption induced by ischaemic stroke. Liposomal transport through the BBB deficits in experimental stroke is mediated by a stepwise impairment of transcellular followed by paracellular barriers. Our data revealed, for the first time, two windows for selective stroke treatment including; a) targeting the NVU in the acute phase to preserve the brain function and minimise the deleterious consequences of stroke, and b) targeting inflammatory cells in the ischaemic brain to shift their polarization towards brain repair. Future studies to test the therapeutic potential of liposomes in ischaemic stroke are warranted to prove their potential utility in accelerating the clinical translation of stroke treatments.

## Materials and Methods

### Materials

Hydrogenated soy phosphatidylcholine (HSPC) and 2-distearoyl-*sn*-glycero-3-phosphoethanolamine-*N*-[methoxy(polyethylene glycol)-2000] (DSPE-PEG_2000_) were kind gifts from Lipoid GmbH (Ludwigshafen, Germany).18:0 PE-DTPA 1,2-distearoyl-*sn*-glycero-3-phosphoethanolamine-N-diethylenetriaminepentaacetic acid (ammonium salt) was purchased from Avanti Polar Lipids (USA). Chloroform and methanol were purchased from Fisher Scientific. Phosphate buffer saline, cholesterol, paraformaldehydes were purchased from Sigma. 1,1’-Dioctadecyl-3,3,3’,3’-Tetramethylindocarbocyanine Perchlorate (DiI) was purchased from Invitrogen Detection Technologies. Polycarbonate extrusion filters (Whatman) 800nm, 200nm, and 100nm were form VWR, UK. PD-10 desalting columns were bought from GE-Healthcare Life Sciences.

### Mice and diets

C57BL/6 male mice (11-12-week-old, weighing 25-30 g; Envigo, UK) were housed in groups of 4–5. All mice were given free access to diet and water and were housed at a constant ambient temperature of 21 ± 2°C and humidity of 40–50%, on a 12-h light, 12-h dark cycle. All experimental procedures using animals were carried out according to the United Kingdom Animals (Scientific Procedures) Act, 1986 and approved by the Home Office and the local Animal Ethical Review Group, University of Manchester and reported in compliance with the ARRIVE guidelines.

### Studies design and exclusion criteria

Calculations of sample sizes were based on power analysis of data from pilot studies and previous experiments. For DiI-Lp accumulation in the brain, we have estimated a mean value of 4.0 E^−5^ for healthy mice and a mean value of 4.3 E^−4^ for MCAo mice with a SD of 1.05 E^−4^. Assuming a significance level of ≤ 0.05 and a power of 80%, the estimated sample size is n=3-4 for detection of 60-70% difference. For the detection of ^111^In-Lp ID in the brain, the mean value for healthy mice is 0.04% of ID and MCAo mice is 0.24% with STD of 0.05% which gives a sample size of n=3-4 mice to detect a deference of 50-60%.

The data were excluded from the analysis in the following conditions; 1) lack of sustained reduction in cerebral blood flow and 2) if signs of subarachnoid haemorrhage or seizure were observed. All experimental procedures using animals were carried out according to the United Kingdom Animals (Scientific Procedures) Act, 1986 and approved by the Home Office and the local Animal Ethical Review Group, University of Manchester and reported in compliance with the ARRIVE guidelines.

### Induction of focal cerebral ischaemia

Focal ischaemic stroke was induced by transient middle cerebral artery occlusion (MCAο) as previously described.[10, 62] Briefly, under 2% isoflurane anaesthesia (in a mixture of 30 % oxygen and 70 % nitrous oxide), the carotid arteries were exposed and a 6–0 silicon rubber-coated monofilament (Doccol, USA) with a 2-mm tip (210μm diameter, coating length 405 mm) was inserted into the left common carotid artery and advanced along the left internal carotid artery 10mm after the left carotid bifurcation. Cerebral blood flow was monitored in all mice by laser-Doppler (Moor Instruments, UK) and MCAo was confirmed by a drop in cerebral blood flow of at least 40-50% of baseline. If this drop in blood flow was not attained, animals were excluded from the analysis. After 20min occlusion, reperfusion was achieved by withdrawing the filament and the wound was sutured. During surgery, core body temperature was monitored using a rectal probe and maintained at 37 ± 0.5°C, using a homoeothermic blanket. Before recovery all mice were given saline (0.5 ml, S.C) and buprenorphine (0.05mg/kg S.C). After surgery, mice were weighed every day and assessed for their general well-being. Body weight data were presented as a % weight change compared with body weight on the day of surgery. Assessment of cerebral ischemia was performed using the a 28-point neurological scoring system[63]. Foot fault test was also performed to confirm ischaemic stroke model as previously described [62]. At the end time point of each group, MCAo mice were placed on an elevated grid surface with grid openings of 2.5cm^2^. During locomotion on the grid, the number of foot slips of both the ipsilateral and contralateral limbs was recorded. Ipsilateral refers the ischaemic side of the body (left) and contralateral limbs are those on the opposite side (right). Tests repeated for three times for each mouse and each trial lasted for 1min. An interval of at least 1min was kept between each trial. The total number of errors of each side is recorded and expressed as % of contralateral foot slips.

### Preparation of DiI-labelled liposomes (DiI-Lp)

DiI-labelled liposomes composed of HSPC:Chol:DSPE-PEG_2000_ 56.3:38.2:5.5 mol/mol % were prepared by thin film hydration method followed by extrusion. Briefly lipids dissolved in chloroform: methanol mixture (4:1) were mixed in round bottom flask and 5mol% of DiI in ethanol (1mg/ml) was added to the lipid mixture. Organic solvents were then evaporated to produce the lipid film.[64–66] Lipid films were kept protected from light and hydration was performed with HBS (20mM HEPES, 150mM NaCl, pH 7.4) to a final lipid concentration of 12.5mM. To produce small unilamellar liposomes, the size was reduced by extrusion though 800nm and 200nm polycarbonate filters 5 times each then 20-40 times through 100nm membranes using a mini-Extruder (Avanti Polar Lipids, Alabaster, AL).

### Preparation and characterisation of ^111^In-labelled liposomes (^111^In-Lp)

To quantify and study the accumulation of liposomes into the brain in real-time SPECT/CT imaging and gamma counting of the liposomes were performed after radiolabelling with radioactive indium (^111^In). Briefly, 25mM (total lipid concentration) of HSPC:Chol:DSPE-PEG_2000_:PE-DTPA 56.3:38.2:5.5:1 mol/mol % liposomes were prepared as described above using thin film hydration method. Hydration of the lipid film was done with freshly prepared ammonium acetate buffer (0.095M, pH 5.5) at 60°C followed by extrusion to reduce the size of the liposomes. Subsequently liposomes were radiolabelled by 1h incubation with radioactive ^111^InCl_3_ (11MBq/2.5µmol lipids) in 2.0M ammonium acetate pH 5.5. Incubation carried out at room temperature with continuous vortexing every 5 minutes. At the end of incubation 0.1M EDTA (1/20 of the total volume) was added to chelate any free ^111^In. To determine the Radiolabelling efficiency, any unbound ^111^In and ^111^In-EDTA were removed with PD-10 column pre-equilibrated with HBS pH 7.4. Aliquots of each final product were diluted five folds in PBS and then 1 μl was spotted on silica gel impregnated glass fibre sheets (PALL Life Sciences, UK). The strips were developed with a mobile phase of 50mM EDTA in 0.1M ammonium acetate and allowed to dry before analysis. This was then developed and the autoradioactivity quantitatively counted using a Cyclone phosphor detector (Packard Biosciences, UK). The immobile spot on the TLC strips indicated the percentage of radiolabelled ^111^In-Lp, while free ^111^In was detected as the mobile spots near the solvent front. Very minimum free ^111^In was detected to yield radiolabelling efficiency of >85% as displayed in Figure S2. The radiolabelling stabilities of the final product of ^111^In-Lp were studied after five times dilution in both 50% serum or PBS and then incubated at 37°C up to 48h. At different time-points (0, 1 and 24h), 1μl of the aliquots was spotted on silica gel impregnated glass fibre sheets and then developed and quantified as described above. No significant release of ^111^InCl_3_ was detected after incubation with PBS and minimum free ^111^InCl_3_ was detected after incubation in 50% for 2 days (Figure S2).

### Liposomes characterization

Liposome size and surface charge were measured using Zetasizer Nano ZS (Malvern, Instruments, UK). Samples were diluted 100 times with purified distilled water before measurements. Triplicates measurements were recorded, and the data expressed as average ± S.D. Fluorescen intensity of DiI-Lp was recorded using Carry Eclipse fluorescence spectrophotometer, (Agilent technology). Samples were first diluted 200 times in HBS and recorded at 518nm/565nm excitation/emission wavelengths (slit 5/10).

### Optical imaging of Lp-DiI accumulation into the brain

Ex vivo optical imaging was used to study the accumulation of DiI-Lp into the brain by detecting the optical fluorescence signal of the liposomes. IVIS imaging was performed shortly after cardiac perfusion with iced-cold 0.9% saline followed by 4% PFA in order to remove any DiI-Lp that are still circulating in the blood. Brain tissues were extracted either 2h or 24h after I.V administration of DiI-Lp into MCAo mice (at 0.5h, 4h, 24h and 48h after 20min MCAo and reperfusion) and healthy mice. Imaging were performed with IVIS Lumina II imaging system (Caliper Life Sciences Corp., Alameda, CA) at 535nm/DsRed excitation and emission filters with 0.5 sec exposure. DiI-Lp total fluorescence intensity in the brain was quantified by drawing a region of interest (ROI) that covers the whole brain and expressed as total efficiency. We would like to emphasise that this method offers ‘semi-quantitative’ estimations of fluorescence intensity of the liposomes in the brain and the absolute quantification of the liposomes in the brain as % of I.D was measured in a separate experiment by gamma counting of ^111^In-liposomes (as explained below).

### Single photon emission computed tomography (SPECT/CT)

Mice were subjected to anaesthesia via the inhalation of 2.5% isoflurane in a mixture of 30 % oxygen and 70 % nitrous oxide. Each animal was then intravenously injected with 200ul of the radioactive ^111^In-Lp (8-9 MBq). At different time points after injection (t= 0-1h, & 24h) SPECT/CT imaging was carried out using a Nano-Scan^®^ SPECT/CT scanner (Mediso, Hungary). SPECT images were obtained in 20 projections over 40-60 min using a 4-head scanner with 1.4 mm pinhole collimators. CT scans were taken at the end of each SPECT acquisition using a semi-circular method with full scan, 480 projections, maximum FOV, 35 kV energy, 300 ms exposure time and 1-4 binning. Acquisitions were done using the Nucline v2.01 (Build 020.0000) software (Mediso, Hungary), while reconstruction of all images and fusion of SPECT with CT images was performed using the Interview™ FUSION bulletin software (Mediso, Hungary). The images were further analysed using VivoQuant 3.0 software (Boston, US) where the SPECT images with scale bars in MBq were corrected for decay and for the slight differences in radioactivity in the injected doses between animals.

### Cranial window implantation and multi-photon imaging

Cranial windows were implanted following the protocol published by Goldey et. al 2014 [61]. Animals were anesthetized with 2.5% isoflurane in 100% room air. After an injection of Metacam (50 µl SC in 1:10 water) and Dexafort (30 µl IM) the scalp was removed over the stroked hemisphere. Then a metal head plate was mounted (Narishige CP-2, Japan) to allow stereotaxic fixation under the two-photon microscope (Leica SP8 MP, UK HC FLUOTAR L 25×0.95 WATER dipping lens). A circular piece of bone with a diameter of 3mm was removed over the somatosensory area of the cortex that is supplied by the MCA. Once the skull was removed a circular cover slip (Warner Instruments, USA) was glued in its place using dental cement (Sun Dental, Japan). After the surgery animals were housed individually and allowed to recover for at least one week and monitored for normal behaviour like nest building and grooming.

*In vivo* multiphoton imaging was carried out under 1.5% isoflurane anaesthesia in a 50:50 mix of oxygen and nitrous oxide at 0.5 hours and 48 hours after MCAo. Once anaesthesia was induced DiI-Lp (100µl), cascade blue dextran 3k (50µl) and Alexa-488 albumin (50µl) were injected via the tail vein. For the imaging itself we selected a stroked area of the brain that displayed different size vessels as well as large areas of parenchyma. 3D stacks were recorded up to a depth of 150µm and z projection images calculated from these with ImageJ.

### Tissue processing

Tissue processing was carried out 2h or 24h after liposomes injections. Mice were terminally anaesthetised with isoflurane and perfused transcardially with iced-cold 0.9% saline and then fixed with 4 % paraformaldehyde (PFA) in 0.1M phosphate buffer saline (PBS). Subsequently, brain samples were removed and post-fixed (in 4% PFA) overnight, cryoprotected in 30% sucrose for 1-2d and snap frozen in isopentane on dry ice. Coronal brain sections (30μm) were cut on a freezing sledge microtome (Bright 8000–001, Bright Instrument Co Ltd, UK) and stored in cryoprotectant (30% ethylene glycol, 20% glycerol in 0.2M phosphate buffer) at −20°C until further processing.

For electron microscopy (EM), mice were perfused with 0.9% saline for 1min at 10ml/min followed by fixative (4% PFA and 2.5% glutaraldehyde in 0.2M HEPES) for 3min. After that brains were removed and post-fixed overnight. Slices were cut at 1mm think then selected areas of striatum and cortex at 0.14mm from the bregma were dissected out and processed for EM as previously described[10]. Briefly, after primary fixation tissues were fixed for 1 h with 1.5% potassium ferrocyanide and 2% osmium tetroxide (weight/vol) in 0.1M cacodylate buffer and 1% uranyl acetate at 4°C overnight. The next day, samples were dehydrated with serial dilutions of alcohol and embedded in TAAB Low viscosity epoxy resin (TAAB, UK). Ultrathin sections (70nm) were cut from resin-embedded samples on an ultramicrotome (Reichert Ultracut), mounted on Formvar-coated grids and viewed on a FEI Tecnai 12 Biotwin Transmission Electron Microscope. Images were acquired with Gatan Orius SC1000 CCD camera.

### Assessment of ischaemic damage

Brain sections were stained with cresyl violet and the infarct volume was calculated by measuring the areas of neuronal loss at eight defined coronal levels as previously described [67, 68]. On each section the area of damage was measured using ImageJ (NIH, Bethesda, MD, USA), adjusted for oedema and the volume of damage calculated by integration of areas of damage with the distance between coronal levels using GraphPad Prism 7, Software. The volume of damage was expressed as the total amount of ischaemic damage. For the assessment of haemorrhagic transformation, haematoxylin and eosin (H&E) staining was performed. The area of red blood cells was measured in the same way as the infract volume was calculated and compared between the groups.

### Assessment of BBB permeability to IgG

To assess BBB permeability, endogenous IgG accumulation into the brain was visualised by peroxidase-based immunohistochemistry. Free-floating serial brain sections (30µm thick) were washed 3 times with PBS and endogenous peroxidase activity and non-specific staining were blocked by 10min incubation in 0.3% H_2_O_2_ followed by washing (3 times in PBS, 10min each). After that 1h blocking with 10% normal horse serum (NHS) in 0.3% Triton X-100 PBS (PBST) was performed before overnight incubation with biotinylated anti mouse IgG (1:250 in in 0.3% PBST, Vector Laboratories) at 4°C. Sections were then incubated with avidin–biotin–peroxidase complex, and colour-developed using a freshly prepared diaminobenzidine (DAB) solution. To ensure comparable DAB staining between sections is achieved, the time of the colour change was recorded, and DAB applied for each subsequent sample for same amount of time.

### Immunohistochemistry

Free-floating serial brain sections (30µm thick) were washed 3 times in PBS for 10min and blocked for 1h in 10% normal goat serum (NGS) in 0.3% PBST. This was followed by an overnight incubation with primary antibody in 2% NGS in PBST at 4°C. The primary antibodies used in the study are explained in details below; Chicken antiGFAP (abcam AB4674, 1:500), Rabbit antiIb1a (Wako 019-19741, 1:500 DF), Rat Anti-mouse CD31 (BD Pharmingen, 550274, 1:100), Mouse anti-mouse Cav1 (BD Biosciences, 1, 610407, 1:50), Mouse anti-mouse NeuN (Millipore, MAB377, 1:100), and Rabbit Anti-MBP (Abcam, ab40390, 1:200). After incubation with the primary antibodies sections were washed 3 times in PBS and incubated with fluorescently labelled secondary antibodies in 2% NGS in PBST. To visualise the primary antibodies, the following secondary antibodies were used; goat anti-Ck Alexa Fluor^®^ 488 conjugate (Invitrogen A11039, 1:500), goat anti-Rabbit Alexa Fluor^®^ 647 conjugate (Invitrogen, A27018 1:200), goat anti-mouse Alexa Fluor^®^ 488 conjugate (Invitrogen A11001, 1:500), goat anti-Rat Alexa Fluor^®^ 647 conjugate (Invitrogen, A-21247, 1:200). At the end samples were washed 3 times in PBS and transferred on non-gelatine coated slides and left to dry overnight before slides were then coverslip with ProLong Gold Antifade Mountant with DAPI (Thermo Fischer Scientific, Inc., USA). Images were collected on either SP5 inverted microscope (for Cav1-1/CD31 & NeuN/MBP) or SP8 inverted microscope (for Iba1/GFAP) using 63x objective in the striatum and the outer cortex at bregma −0.58 mm. Total microglia were counted and the percentage of DiI-Lp positive microglia recorded. The activation state of microglia was scored on activation scale of 0-3 based on their morphologies using a scoring system described before [69].

### Gamma scintigraphy

For a quantitative assessment of ^111^In-Lp in the brain, a cut and count method was used. Mice were anaesthetized by isofluorane inhalation and each mouse was injected *via* the tail vein with 200μl containing ^111^In-Lp labelled with approximately 8-9 MBq. 24h after injection, mice were perfused with iced cold saline (0.9%) followed by PFA (4%) to remove any ^111^In-Lp from the blood before brain tissues were collected. Each sample was weighted and counted on a gamma Counter (Perkin Elmer, USA), together with a dilution of the injected dose with dead time limit below 60%. The results were represented as the percentage of the injected dose (% ID/gm tissue ± SEM), n=4-5 mice per group.

### Data and statistical analyses

Statistical analysis of the data was performed using Graph Pad Prism 7 software. Two-tailed unpaired student t-test and one-way analysis of variance followed by the Tukey multiple comparison test were used and p values < 0.05 were considered significant. For all analyses, data are represented as mean ± standard error of the mean (SEM), unless otherwise indicated.

## Acknowledgments

The authors acknowledge Lipoid Co. (Germany) for the lipid sample gifts, the staff at the bioimaging facility at the University of Manchester for advice on confocal imaging and data analysis, and Aleksandr Mironov of the Faculty of Life Sciences EM Facility for his assistance with TEM experiment. They also wish to thank the Biological Services Facility at the University of Manchester for expert animal husbandry.

## Author Contributions

ZA initiated, designed, planned, and led the study, performed almost all of the experimental work and data analysis, and drafted the manuscript. DJ performed SPECT/CT imaging and radiolabelling stability assays. SA Helped in the initial stages of the project with data analysis and IHC experiments. RY provided training on MCAo model and gave advice on the stroke model and data analysis through out the project. MH performed CSF samples collections and provided continuous advice on data analysis and interpretation. GC provided technical training on histological staining and cardiac perfusion. IS performed cranial window implantation and multiphoton imaging and data analysis. SA provided continuous guidance in the conceptual design of the work and data interpretation, reviewed and edited the manuscript. KK conceptualized the study, designed, planned, discussed the findings, reviewed, edited the manuscript, and overall supervised the work.

## Supplementary Materials

**Figure S1:** Physicochemical characterisation of DiI-Lp.

**Figure S2:** Characterisation of ^111^In-Lp.

**Figure S3:** Selective liposomal accumulation into ischaemic cerebral cortex.

**Figure S4:** Selective liposomal accumulation into the ipsilateral striatum 24h after I.V administration.

**Figure S5:** Confocal images of Cav-1 expression and DiI-Lp accumulation in contralateral striatum and contralateral cortex (2h after DiI-Lp I.V administration).

**Figure S6:** Confocal images of Cav-1 expression and DiI-Lp accumulation in contralateral striatum and contralateral cortex (24h after DiI-Lp I.V administration).

**Figure S7:** Confocal images of DiI-Lp co-localisation with neurons and oligodendrocytes in the ipsilateral striatum.

**Figure S8:** Confocal images of DiI-Lp co-localisation with neurons and oligodendrocytes in the ipsilateral cortex.

