## Supplementary material for "Selective Liposomal Transport Through Blood Brain Barrier Disruption in Ischaemic Stroke Reveals Two Distinct Therapeutic Opportunities": Figure 1 to 8

**Affiliations:**

### Supplementary Materials

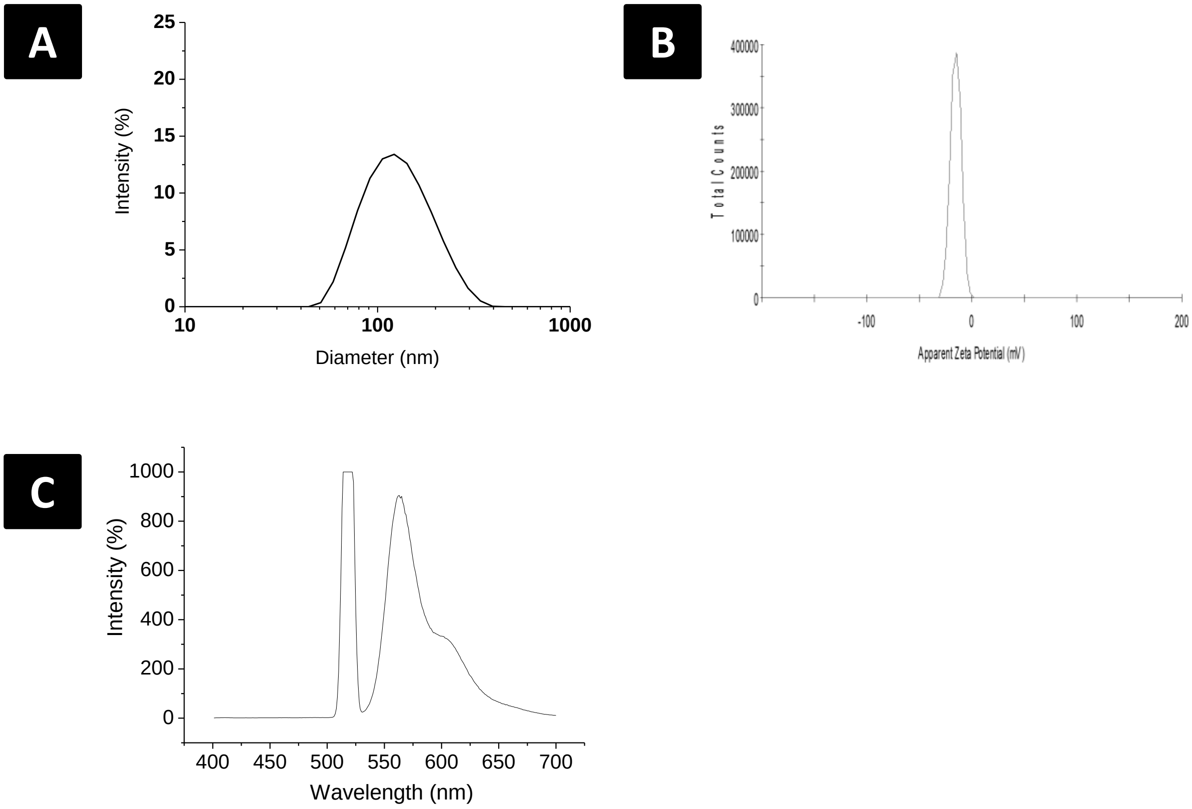


**Figure S1:** Physicochemical characterisation of DiI-Lp showing; (A) hydrodynamic diameter. (B) zeta potential and (C) fluorescent intensity at excitation wavelength of 518nm and emission wavelength 565 nm.

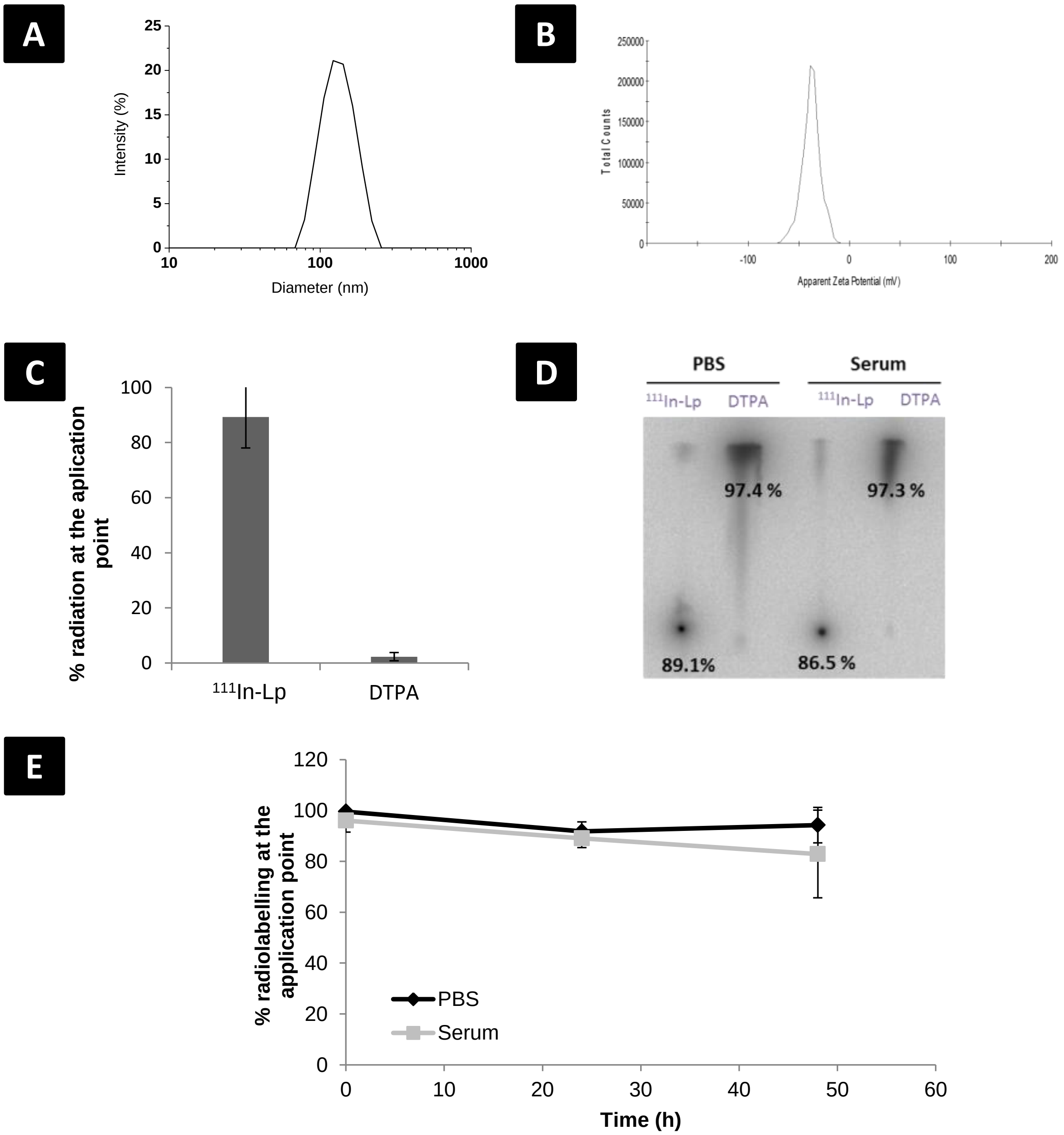


**Figure S2:** Characterisation of $^{111}$In-Lp showing; (A) hydrodynamic diameter. (B) zeta potential, (C&D) radiolabelling efficiency and (E) radiolabelling stability overtime in PBS and 50% serum.

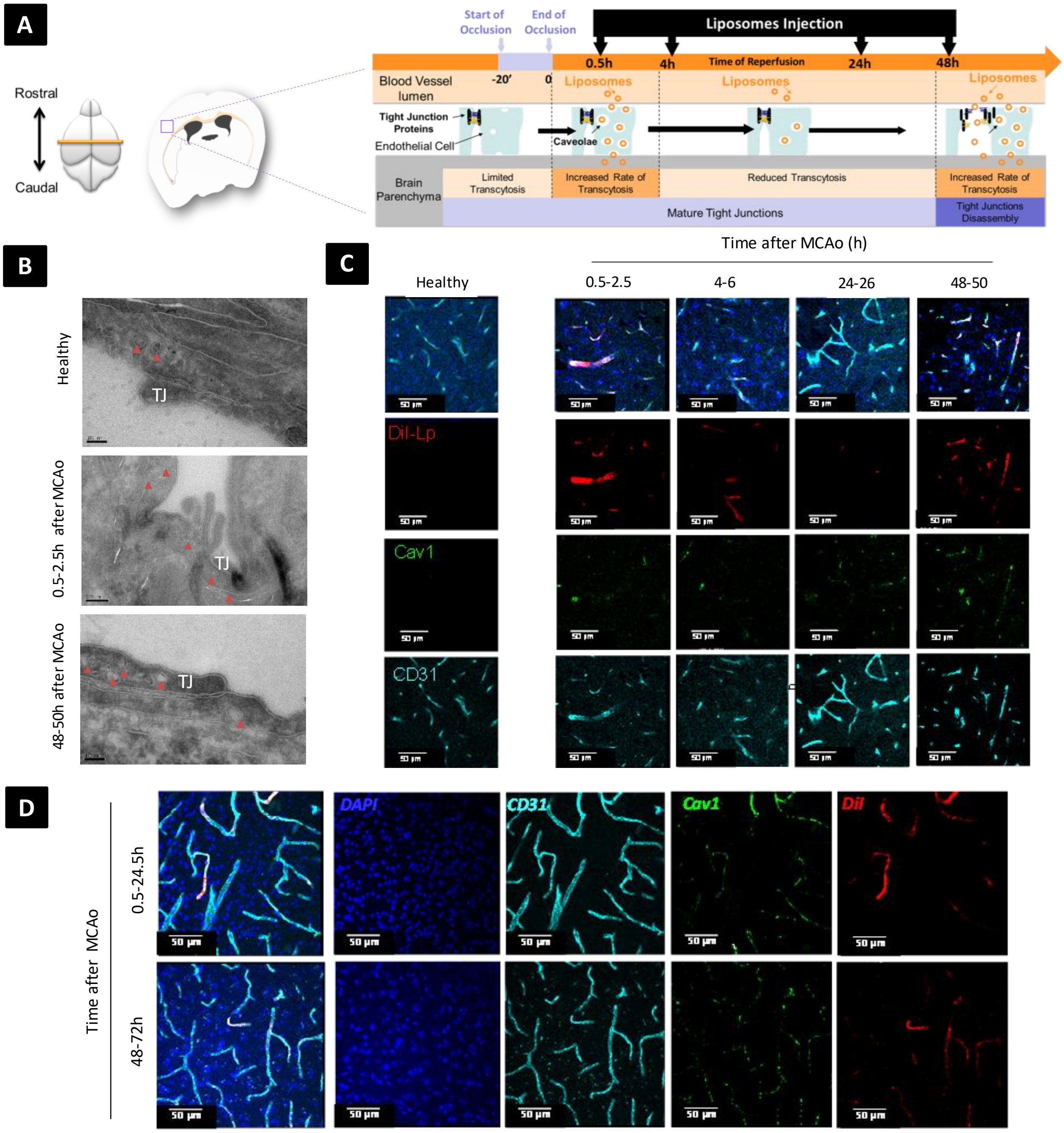


**Figure S3: Selective liposomal accumulation into ischaemic cerebral cortex**. (A) Schematic presentation of the experimental plan and the time frame for DiI-Lp intravenous administration after 20min of MCAo. (B) Ultrastructural changes in the brain endothelial cells in the cerebral cortex studied by EM early and late after MCAo and compared to naïve healthy brain. (C&D) Immunofluorescent staining showed co-localisation of DiI-Lp leakage with areas of enhanced Cav-1 expression when assessed both 2h and 24h after DiI-Lp I.V administration. Maximum DiI-Lp leakage detected when administered at 0.5h or 48h after MCAo indicating a biphasic window of therapeutic opportunities.

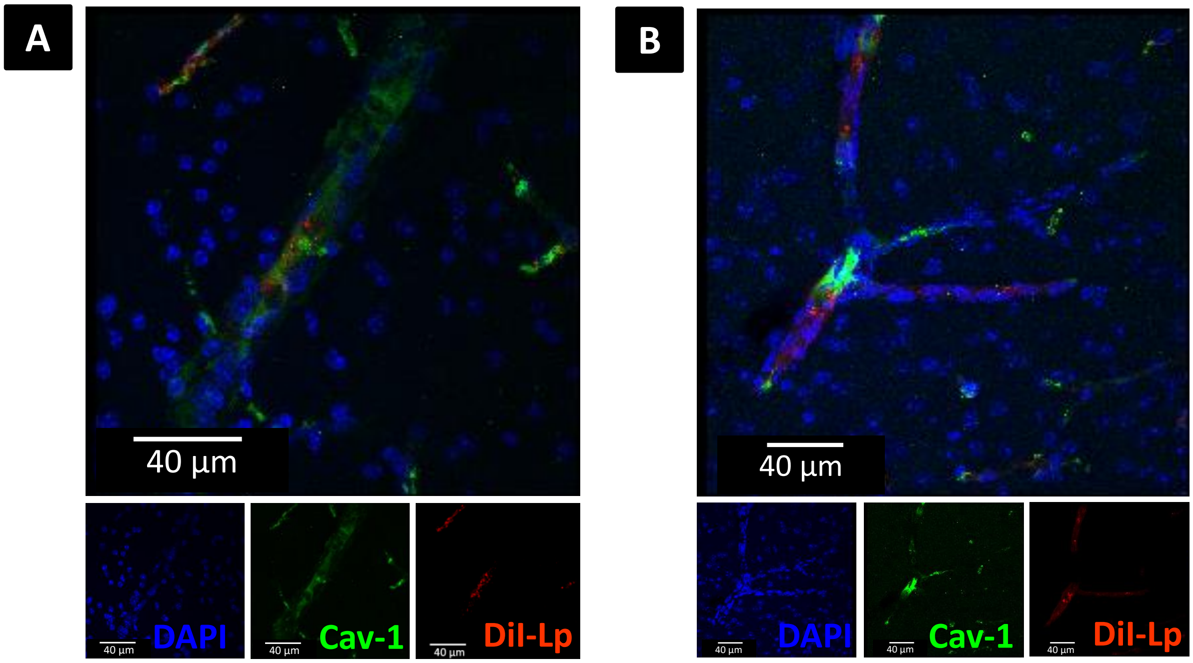


**Figure S4: Selective liposomal accumulation into the ipsilateral striatum24h after DiI-Lp I.V administraion.** Representative high magnification (100x with zoom factor 2) immunofluorescecet imaging of ipsilateral striatum at -0.58mm bregma showing DiI-Lp signal (red) inside Cav-1 positive vesicles of the brain endothelial cells. Confocal images were obtained 24h after I.V administration of DiI-Lp into MCAo mice at; (A) 0.5h and (B) 48h following reperfusion.

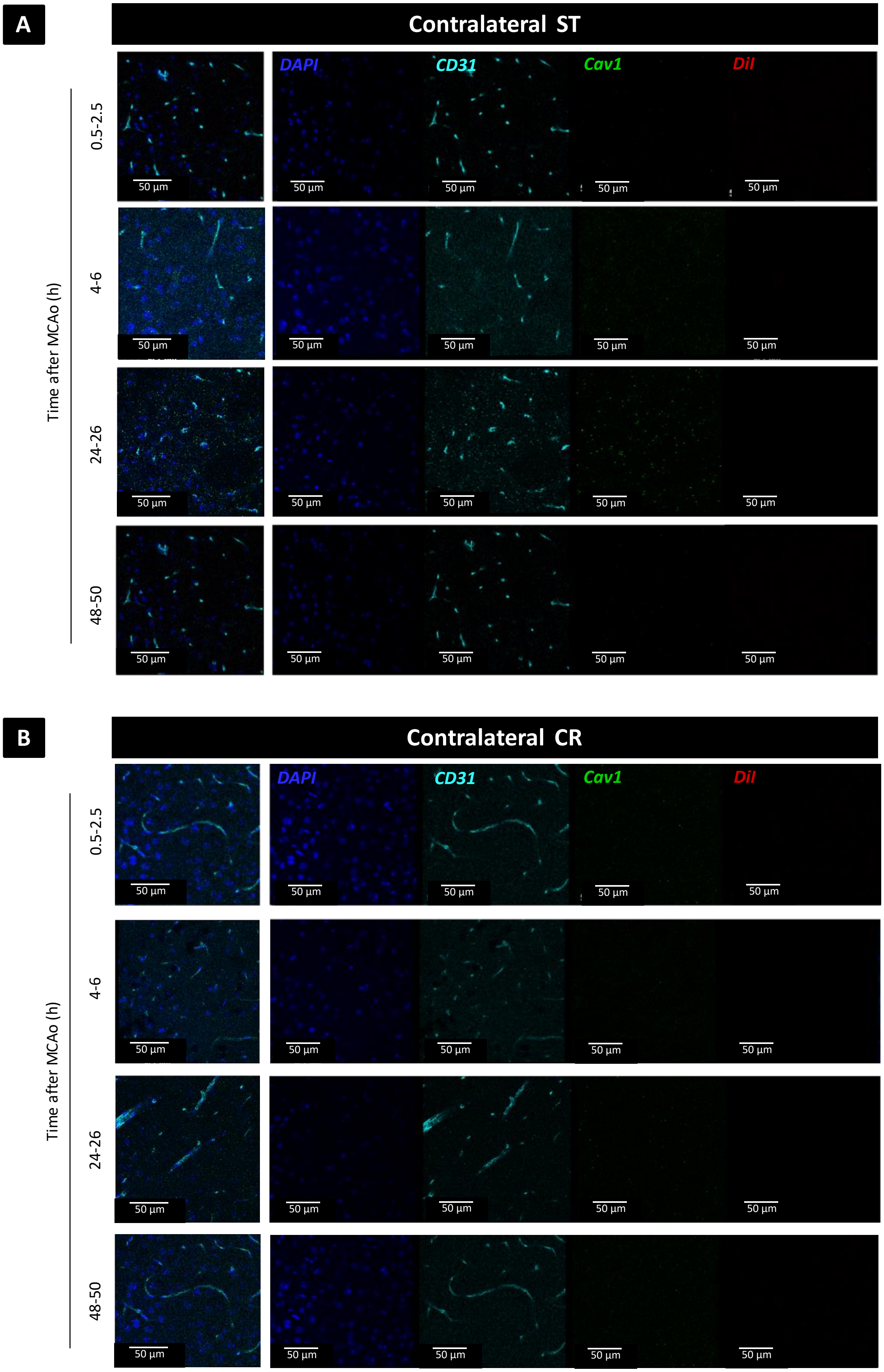
A
Contralateral ST
DAPI
CD31
Cav1
DiI
Time after MCAo (h)
0.5-2.5
4-6
24-26
48-50
50 µm
B
Contralateral CR
DAPI
CD31
Cav1
DiI
Time after MCAo (h)
0.5-2.5
4-6
24-26
48-50
50 µm

**Figure S5:** Confocal images of (A) contralateral striatum (ST) and (B) contralateral cortex (CR) indicated minimum expression of Cav-1 and no DiI-Lp accumulation which confirm the selective accumulation of liposomes into the brain after ischaemic stroke. Images were obtained 2h following DiI-Lp I.V administration.

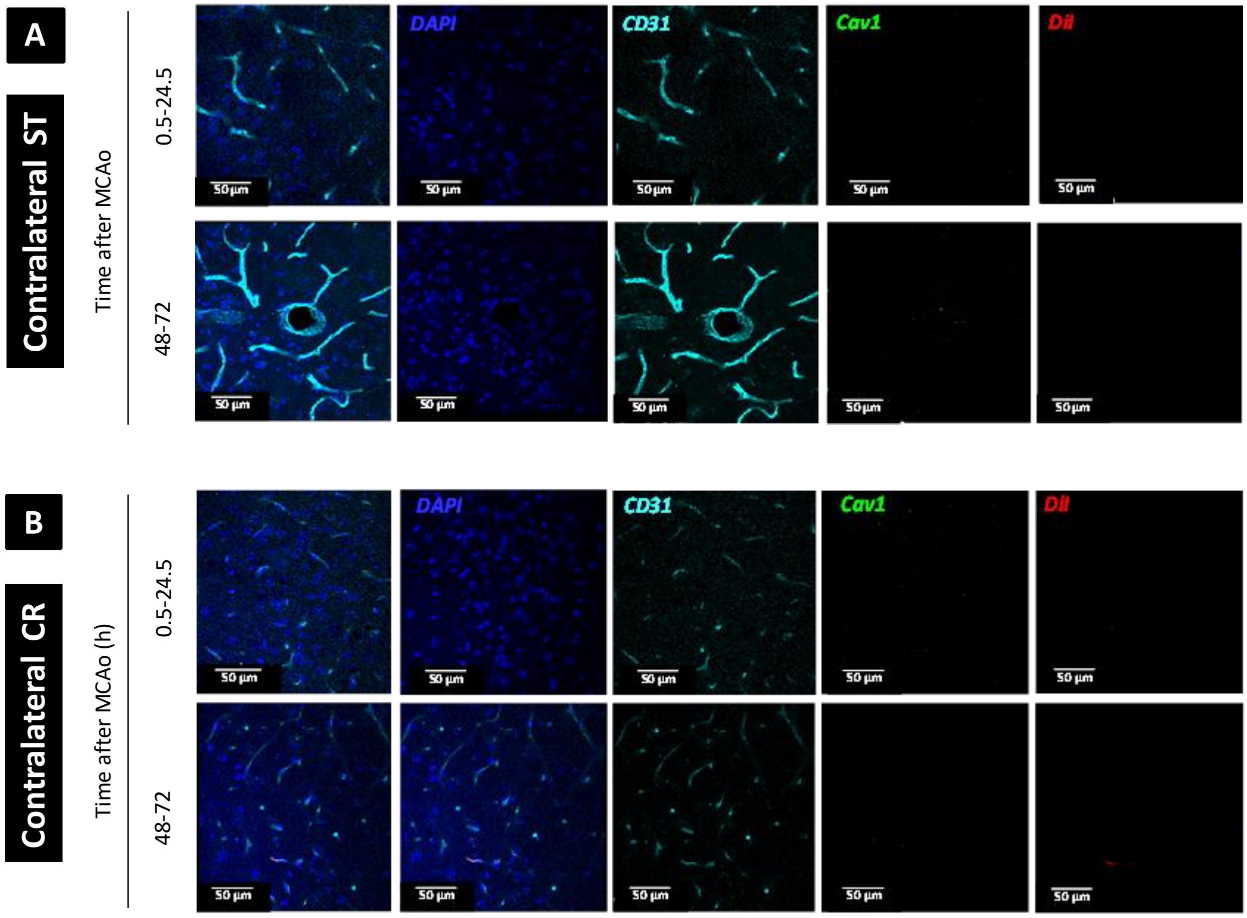


**Figure S6:** Confocal images of (A) contralateral striatum (ST) and (B) contralateral cortex (CR) 24h after DiI-Lp I.V administration into MCAo mice at 0.5h or 48h after reperfusion Images indicated limited Cav-1 expression and no DiI-Lp accumulation.

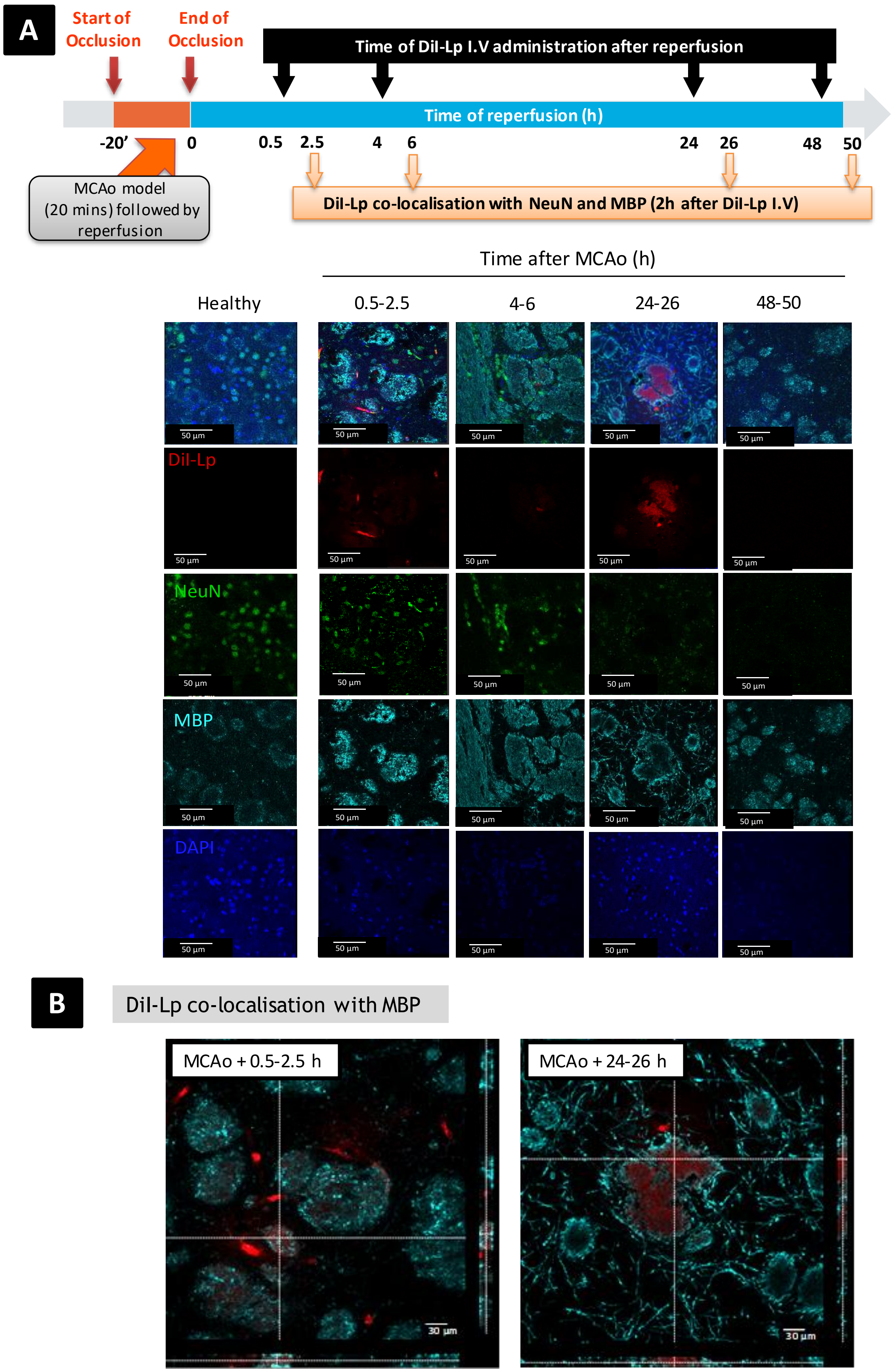
A
Start of Occlusion
End of Occlusion
Time of DiI-Lp I.V administration after reperfusion
Time of reperfusion (h)
-20'
0
0.5
2.5
4
6
24
26
48
50
MCAo model (20 mins) followed by reperfusion
DiI-Lp co-localisation with NeuN and MBP (2h after DiI-Lp I.V)
Time after MCAo (h)
Healthy
0.5-2.5
4-6
24-26
48-50
DiI-Lp
NeuN
MBP
DAPI
50 µm
B
DiI-Lp co-localisation with MBP
MCAo + 0.5-2.5 h
MCAo + 24-26 h
30 µm

**Figure S7:** Confocal images of DiI-Lp co-localisation with Neurons and oligodendrocytes in the ipsilateral striatum. A) 2h following DiI-Lp I.V administration into MCAo mice indicating no clear co-localisation of DiI-Lp with Neurons (green) and occasional co-localisation with oligodendrocytes (Cyan). B) Representative images of DiI-Lp co-localisation with MBP marker.

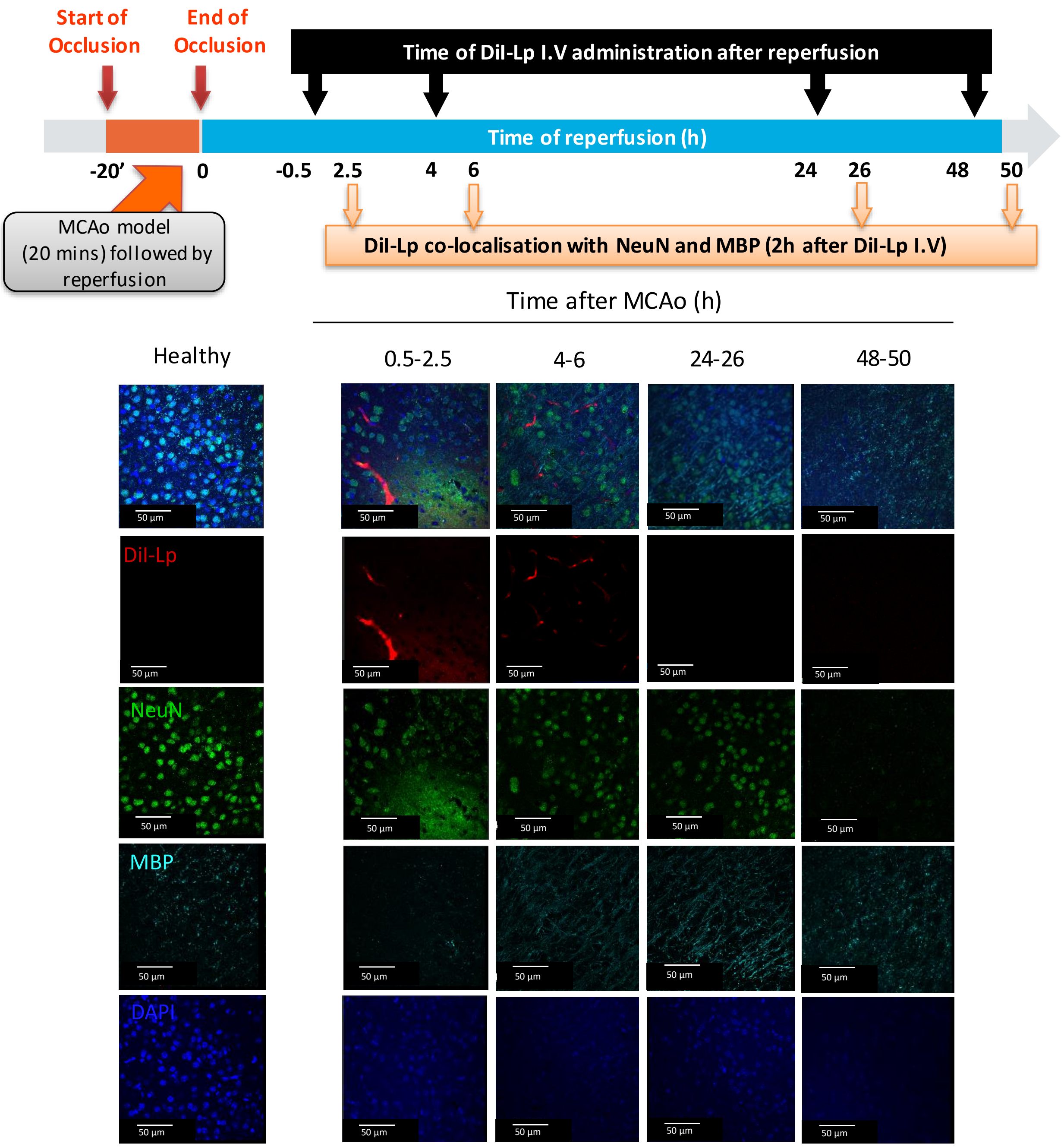


**Figure S8:** Confocal images of DiI-Lp co-localisation with Neurons and oligodendrocytes in the ipsilateral cortex 2h following DiI-Lp I.V administration into MCAo mice indicating no clear co-localisation of DiI-Lp with Neurons (green) and oligodendrocytes (Cyan).
